# Three-dimensional redistribution of pelagic fish aggregations associated with floating offshore wind farms

**DOI:** 10.64898/2026.01.17.700020

**Authors:** Shimpei Tsuchida, Shingo Fujimoto, Siti Syazwani Azmi, Miyuki Hirose, Katsuya Hirasaka, Yukuto Sato, Mitsuharu Yagi

## Abstract

Floating offshore wind farms (F-OWFs) are rapidly expanding into offshore pelagic environments, yet their ecological consequences remain poorly resolved because quantitative, non-invasive monitoring is challenging at sea. Here, we integrated environmental DNA (eDNA) metabarcoding and scientific echosounding to assess fish community composition and three-dimensional aggregation structure around a commercial-scale F-OWF off the Goto Islands, Japan. We compared four stations adjacent to turbines with four offshore control stations under comparable environmental conditions across five sampling periods (April and August 2024; May, June, and August 2025) and three depth layers (5, 50, and 80–160 m). eDNA metabarcoding detected 126 fish species, and community structure based on the 30 most frequently detected taxa did not differ significantly between the F-OWF and control areas (Bray–Curtis PERMANOVA, p = 0.60). Species richness varied strongly with sampling period and water layer, with a significant water layer x period interaction, whereas overall richness did not differ between areas (Mann–Whitney U test, p = 0.18). In contrast, acoustic surveys revealed a marked difference in vertical structuring of fish aggregations: in the control area, NASC per mile increased with depth, while this depth-dependence disappeared near turbines. Moreover, the weighted mean normalized depth (WMND) indicated a shallower vertical center of aggregation in the F-OWF area (0.34) than in the control area (0.24), consistent with turbine-associated redistribution in the water column. Although voyage-level integrated NASC per mile did not differ significantly between areas (t-test, p = 0.83), mean values were higher near turbines. Together, these results indicate that F-OWFs can be associated with changes in the three-dimensional organization of fish aggregations without producing pronounced shifts in dominant taxonomic composition. Our study demonstrates the value of combining eDNA metabarcoding and acoustics for evaluating ecological effects of offshore renewable-energy infrastructures and provides a framework for standardized, long-term monitoring as F-OWF deployment accelerates globally.

## INTRODUCTION

Recent global efforts to mitigate climate change have intensified the demand for decarbonization, leading to a rapid expansion of renewable energy technologies. Among these, offshore wind farms (OWFs) have gained particular prominence because they can harness strong and stable wind resources at sea. According to the Global Wind Energy Council, an additional 8 GW of offshore wind capacity was installed worldwide in 2024, bringing the total to 83 GW—an increase of approximately 11% from the previous year (GWEC, 2025). Although early OWF developments were concentrated in shallow coastal waters of Europe, the saturation of suitable nearshore sites and the pursuit of more powerful wind regimes have driven installations progressively farther offshore and into deeper waters (Díaz and Guedes Soares, 2020). Reflecting this global trend, large-scale OWF deployment is now underway across many regions, including Asia and North America.

Bottom-fixed offshore wind farms (B-OWFs) currently represent the dominant form of offshore wind development, with monopile foundations being the most widely adopted design. This system involves driving large steel piles vertically into the seabed to support turbine structures (Chen and Kim, 2022). The construction, operation, and maintenance of B-OWFs can introduce various anthropogenic stresses into the marine environment, including underwater noise and electromagnetic fields (Kazama, 2012). In particular, pile-driving during installation generates exceptionally high-intensity impulsive noise capable of altering the behavior of marine organisms (Thomsen et al., 2006), and such extreme acoustic levels have been widely documented (Norro et al., 2013). Nevertheless, B-OWFs may also exert positive ecological effects once operational. Their foundations can function as artificial reefs that attract fish and other marine organisms (Wilson, 2007), and vessel exclusion zones established under the United Nations Convention on the Law of the Sea (UNCLOS) effectively create de facto no-fishing areas around turbines (United Nations, 1982; Bonsu et al., 2024; Reckhaus, 2022). These protected zones may enhance local biodiversity (Hammar et al., 2016) and benefit commercially important species (Bailey et al., 2014). Recent acoustic studies have further shown elevated fish densities around wind turbine foundations compared with surrounding waters, suggesting localized aggregation effects (Hamana et al., 2025).

A second major class of offshore wind development is floating offshore wind farms (F-OWFs; Fig. 1). The F-OWFs deployed in the present study area use a hybrid-spar configuration comprising a concrete lower section and a steel upper structure stabilized by three mooring chains anchored to the seabed (Sato and Matsunobu, 2021). This design enables installation in much deeper waters than is feasible for bottom-fixed systems, thus greatly expanding potential deployment areas and supporting expectations for rapid global growth. However, because F-OWFs are technically more challenging to construct and remain relatively few worldwide, their ecological impacts are still poorly understood (Farr et al., 2021; Rezaei et al., 2023). Given that floating platforms occupy the upper water column, they may function analogously to fish aggregating devices (FADs) and attract pelagic species. Indeed, a year-long eDNA survey conducted off the Goto Islands demonstrated that F-OWFs exerted a measurable aggregation effect on Japanese jack mackerel (Trachurus japonicus) (Tsuchida et al., 2025). Yet this study focused on a single species, leaving the broader effects of F-OWFs on entire fish communities largely unresolved.

**Fig. 1.**
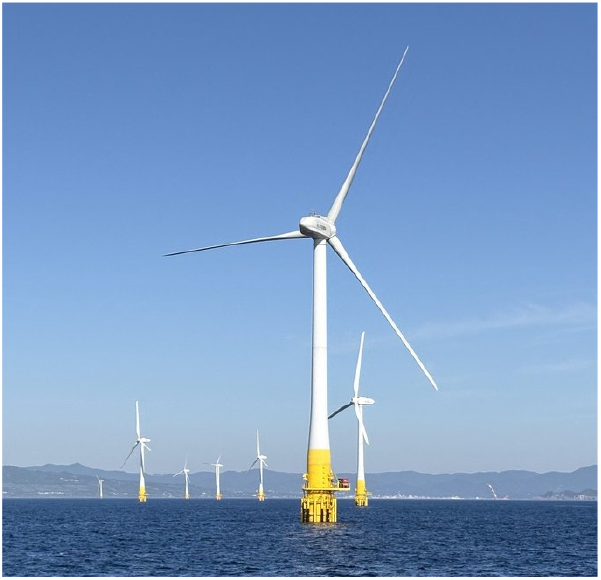
Floating offshore wind turbines (F-OWFs) off the Goto Islands, Japan.

To understand how F-OWFs influence surrounding fish communities, it is essential to clarify both (i) the taxonomic composition of the assemblage and (ii) its three-dimensional spatial structure. Quantitative echosounders provide high-resolution measurements of fish biomass distribution across the water column (Ressler et al., 2012), whereas environmental DNA (eDNA) metabarcoding allows comprehensive and non-invasive assessment of species composition (Miya, 2022). Integrating these complementary approaches offers a powerful framework for evaluating how artificial offshore structures modify the three-dimensional organization of marine ecosystems with unprecedented detail. In this study, we combined acoustic surveys and eDNA metabarcoding to assess fish community composition and vertical distribution around the F-OWFs deployed off the Goto Islands. By comparing turbine sites with nearby control sites, we aimed to disentangle the influence of F-OWFs on local fish assemblages. This integrated approach provides a multidimensional assessment of F-OWF effects and yields new insights into the ecological functions that such offshore infrastructures may introduce into pelagic ecosystems.

## MATERIALS AND METHODS

### Experimental Design

To evaluate spatial patterns in fish community composition and aggregation associated with F-OWFs, we implemented a field-based comparative study design analogous to a treatment–control framework commonly used in ecological assessments. In this context, “treatment” stations were defined as locations situated in the immediate vicinity of F-OWFs, where physical structures and associated ecological influences were expected to be most pronounced, whereas “control” stations were established in areas without turbines and selected to represent comparable offshore environmental conditions.

Specifically, four stations (E1–E4) were positioned around the F-OWFs located approximately 5 km offshore of Fukue Island, Nagasaki Prefecture, Japan (Fig. 2). Four control stations (E5–E8) were established approximately 4 nautical miles (7.4 km) south of the turbine array (Fig. 2). Control stations were selected to closely match the F-OWF stations in terms of water depth, offshore setting, and exposure to regional oceanographic conditions, while being sufficiently distant from the turbine array to minimize direct structural, acoustic, or hydrodynamic influences of the F-OWFs. Although it is not possible to completely exclude all indirect influences in open marine systems, this spatial separation was considered adequate to distinguish areas directly associated with F-OWFs from those representing baseline offshore conditions (Tsuchida et al., 2025).

**Fig. 2.**
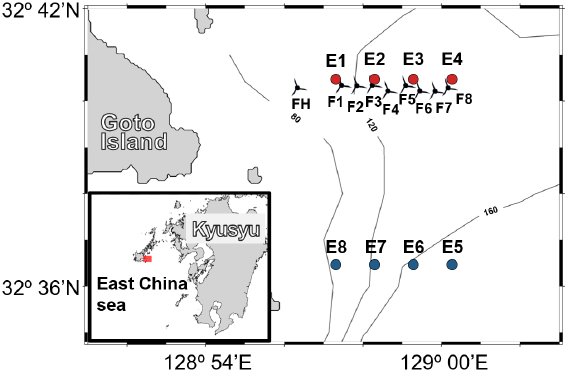
Geographic locations of the eight sampling stations (E1–E8) used for eDNA metabarcoding. Stations E1–E4 correspond to the F-OWF area, whereas E5–E8 represent the control area. The positions of floating offshore wind turbines (FH, F1–F8) are also shown. Coordinates and water depths for all stations and turbines are provided in Supplementary Table S1.

Surveys were conducted aboard the training vessel Kakuyo-maru (155 gross tonnage), Nagasaki University, during 2024 and 2025. At each station, seawater temperature (°C) and salinity were measured using a CTD profiler (SBE-911 plus, Sea-Bird Electronics, Bellevue, WA, USA) to characterize the physical environment and confirm broad comparability between F-OWF and control areas. The F-OWF facility comprised the initial demonstration turbine “Haenkaze” (FH), installed in 2015, and additional turbines that became operational during the study period. By May 2025, seven turbines (FH–F6) were operational; this number increased to eight in June (FH–F7) and nine in August (FH–F8). Each turbine had a rotor diameter of 80 m, a total structure height of 172 m (76 m submerged depth, 56 m hub height), a maximum floater diameter of 7.8 m, and a rated output of 2,100 kW. The geographic coordinates and water depths of all sampling stations and turbine locations are provided in Supplementary Table S1.

Because the number of operational turbines increased over the course of the study, the F-OWF area was treated as a dynamic infrastructure system rather than a static experimental unit. All surveys were therefore interpreted as snapshots of fish community structure under progressively expanding F-OWF deployment, rather than as a before–after impact assessment. This approach precludes strict causal inference but allows for the detection of consistent spatial patterns in fish community composition and aggregation associated with the presence of floating offshore wind infrastructure under comparable offshore conditions.

The experimental design was intentionally structured to integrate two complementary observational approaches—environmental DNA (eDNA) metabarcoding and in situ acoustic surveys—conducted at the same stations and during the same sampling periods. eDNA metabarcoding provided high-resolution information on species composition across multiple depth layers, whereas acoustic surveys quantified the horizontal and vertical distribution of fish aggregations in the water column. By combining these methods within a unified spatial framework, the study aimed to identify whether consistent differences in taxonomic composition and three-dimensional spatial organization of fish communities were associated with proximity to F-OWFs, beyond what would be expected from natural offshore variability alone.

### Meta-barcording

Environmental DNA (eDNA) sampling was conducted in April and August 2024 and in May, June, and August 2025. At each station (Fig. 2), seawater was collected from three depth layers: the surface (5 m), mid-water (50 m), and bottom layer (80–160 m). A total of 3 L of seawater was collected per layer using Niskin bottles attached to a CTD profiler. During each CTD cast, in situ seawater temperature and salinity were simultaneously recorded to characterize the physical environment at each sampling depth (Supplementary Table S2). To prevent contamination, Niskin bottles and sampling hoses were rinsed with chlorine-based detergent before use, and powder-free nitrile gloves were worn during all procedures. A field blank consisting of 3 L of artificial seawater was collected following the same protocol. Immediately after sampling, 3 mL of 10% benzalkonium chloride solution (final concentration 0.01%) was added to each sample to inhibit DNA degradation (Minamoto et al., 2021), and samples were stored in a cool and dark environment until filtration.

Filtration was performed within 48 h after sampling using an aspirator (GAS-1N, AS ONE, Osaka, Japan). During filtration, the apparatus was covered with aluminum foil, and all components were decontaminated with 0.1% sodium hypochlorite between samples. Seawater was filtered through 47-mm-diameter glass fiber filters (Whatman GF/F, pore size 0.7 μm). Filters were wrapped in aluminum foil and stored at −20°C until DNA extraction.

eDNA was extracted using the DNeasy Blood & Tissue Kit (Qiagen, Hilden, Germany) following a modified spin-column protocol based on Ono et al. (2023, 2024) and Yamamoto et al. (2016), as adopted by Tsuchida et al. (2025). Field blank samples were processed in the same manner as environmental samples.

The first PCR (1st PCR) was conducted using MiFish-U universal primers following Miya and Sado (2024). Each 12.0 µL reaction consisted of 6.0 µL KAPA HiFi HS ReadyMix, 2.8 µL of 20-fold diluted primers, 1.2 µL DNase/RNase-free water, and 2.0 µL template DNA. To minimize PCR dropouts, four replicate PCRs were performed per sample. PCR blanks were prepared using DNase/RNase-free water instead of template DNA.

MiFish-U-F: ACACTCTTTCCCTACACGACGCTCTTCCGATCTNNNNNNGTCGGTAAAACTCGTGCCAGC

MiFish-U-R: GTGACTGGAGTTCAGACGTGTGCTCTTCCGATCTNNNNNNCATAGTGGGGTATCTAATCCCAGTTTG

Thermal cycling conditions were: 95°C for 180 s; 35 cycles of 98°C for 20 s, 65°C for 15 s, and 72°C for 15 s; and a final extension at 72°C for 300 s. PCR products were stored at −20°C. First PCR products were purified using AMPure XP magnetic beads (Beckman Coulter). For each reaction, 20 µL of PCR product was mixed with 0.9 µL of magnetic beads, washed twice with 70% ethanol, and eluted in RNase/DNase-free water.

The second PCR (2nd PCR) followed Miya and Sado (2024). Each 15.0 µL reaction consisted of 7.5 µL KAPA HiFi HS ReadyMix, 0.88 µL each of Forward and Reverse primer, 3.88 µL DNase/RNase-free water, and 1.86 µL of purified 1st PCR product. Unique dual indices (UDIs) were added using the Multiplex Oligos for Illumina kit (96 Unique Dual Index Primer Pairs, E6440 and E6442; New England Biolabs), assigning 8-bp index sequences to both i5 and i7 primers. Thermal cycling conditions were as follows: 95°C for 180 s; 98°C for 20 s; 72°C for 15 s (annealing/extension); followed by a final extension at 72°C for 300 s. After amplification, products were verified by agarose gel electrophoresis. Bands within the expected range (275–370 bp) were excised to remove primer dimers and non-specific fragments (<100 bp). Gel-purified products were further cleaned using AMPure XP beads, after which DNA concentration was measured.

Purified libraries were submitted to Macrogen Japan (Tokyo, Japan) and sequenced on the *NovaSeq X* Plus platform using a 2 × 150 bp paired-end configuration on a 10B flow cell. The sequencing generated an average of 4.10 × 10^6^ paired reads per sample. Sequencing data were processed following a standard metabarcoding workflow. Low-quality reads were first removed using *fastp*, after which primer sequences were trimmed from both ends using cutadapt. The remaining forward and reverse reads were merged into single contigs using *vsearch*, and merged sequences were subsequently aggregated for each sample. Unique sequence variants were then compared against the MiFish reference database (ver. 4.07; Miya et al., 2015) using local BLAST searches, and taxonomic assignments were made using an E-value cutoff of 1e^−5^.

### in-situ acoustic survey

Acoustic surveys using a scientific echosounder were conducted in May, June, and August 2025 to quantify the horizontal and vertical distribution of fish aggregations in the F-OWF area and the control area. All acoustic surveys were conducted immediately after eDNA water sampling at the corresponding stations. Surveys were performed using a SIMRAD EK80 echosounder operated from the vessel. The vessel maintained a constant engine setting of 950 rpm during transects, corresponding to an approximate cruising speed of 8 knots. In the F-OWF area, zigzag transects were conducted between the F-OWFs FH and F1–F8, whereas in the control area, transects were run between the points S1–S6 (Fig. 3). Transects from FH toward F8 were designated as Navigation A, and those from F8 toward FH as Navigation B. Similarly, transects from S1 toward S6 and from S6 toward S1 were designated as Navigation C and Navigation D, respectively.

**Fig. 3.**
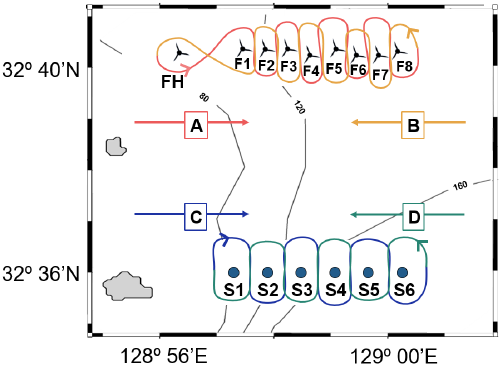
Schematic illustration of zigzag transects conducted for the acoustic survey to characterize the three-dimensional distribution of fish aggregations. Transects in the F-OWF area were conducted between turbine foundations FH–F8, while those in the control area were run between reference points S1–S6. Coordinates and water depths for all stations are provided in Supplementary Table S1.

Acoustic data were collected using a hull-mounted 200-kHz transducer (beam width 7°) operated in continuous wave (CW) mode. The transmitted pulse had a power of 150 W, a pulse duration of 1.024 ms, and a ping interval of 1.0 s. Raw acoustic backscatter data were processed using Echoview software (ver. 9.0; Echoview Software Pty Ltd, Hobart, Australia). A bottom detection line was defined to separate the seafloor from the water column, and surface exclusion zones were set to remove noise caused by bubbles and surface turbulence. Additional noise spikes or anomalous backscatter near the seabed were manually excluded during data cleaning. Acoustic data obtained during Navigation A in May 2025 were omitted from subsequent analyses due to instrument malfunction resulting in excessive noise.

### Data analysis

Fish community composition was assessed based on taxa detected by eDNA metabarcoding. To establish a standardized reference framework for taxonomic assignment, we first compiled a comprehensive list of Japanese marine fish species registered in FishBase (Froese and Pauly, 2025), comprising approximately 4,000 species. Species information was retrieved using the *rfishbase* package (Boettiger et al., 2012) by selecting taxa that met the following criteria: occurrence in Japan (C_Code = 392), native or endemic status, and saltwater habitat designation. Sequence variants obtained from each eDNA sample were taxonomically assigned using this reference framework, and species detection lists were generated for subsequent analyses. For community-level analyses, sequencing reads were aggregated at the species level, and species were retained in the analytical dataset based on their detection across samples.

To test whether fish species richness differed between areas associated with F-OWF and control areas, species richness was calculated for each sample as the total number of detected fish species. Species richness was compared using data pooled across sampling seasons and depth layers to evaluate overall spatial differences while minimizing the influence of short-term temporal variability. Because species count data deviated significantly from normality (Shapiro–Wilk test: W = 0.87, p = 5.35 × 10^−9^), differences between F-OWF and control areas were assessed using a nonparametric Mann–Whitney U test.

To identify factors contributing to variation in fish species richness, we further applied generalized linear models (GLMs). The response variable was species richness per sample, while explanatory variables included water layer (surface, mid-water, bottom), sampling period (April 2024; August 2024; May, June, and August 2025), and area type (F-OWF vs. control). Because the number of operational turbines increased progressively after January 2025, potentially introducing temporal heterogeneity in turbine-related effects, interaction terms between sampling period and water layer were included in the model. Species richness data exhibited overdispersion; therefore, a negative binomial GLM (GLM-NB) was employed. The overall significance of explanatory variables was evaluated using Type II analysis of deviance.

To visualize species composition and relative dominance patterns at each station, stacked bar charts were constructed based on the relative contribution of detected fish taxa in the F-OWF and control areas. To statistically evaluate whether fish community structure differed between areas, permutational multivariate analysis of variance (PERMANOVA; 999 permutations) was conducted based on Bray–Curtis dissimilarity. To reduce the influence of sequencing depth variability and to focus on ecologically dominant taxa, analyses were restricted to the 30 species (Supplementary Table S3) with the highest total read counts across all samples. For each sample, read counts of these 30 species were normalized so that their sum equaled one, yielding relative dominance values used for multivariate analyses.

Differences in fish aggregation density and spatial distribution between the F-OWF and control areas were assessed using acoustic survey data. Fish biomass was quantified using the nautical area scattering coefficient (NASC), a widely applied acoustic proxy for fish density (Dornan et al., 2019; Lancaster et al., 2025). Acoustic backscatter was integrated into spatial bins of 0.5 nautical miles horizontally and 5 m vertically. To isolate fish-derived signals, an integration threshold of −70 dB re 1 m^−1^ was applied, and values below this threshold were excluded from analyses (De Robertis et al., 2019). To account for differences in transect length among surveys, NASC values were standardized by navigation distance (miles), yielding NASC per mile. To minimize contamination from non-biological signals, a bottom exclusion line was set 3 m above the detected seabed to remove bottom backscatter (Jones et al., 2011), and a surface exclusion line was set at 10 m depth to eliminate near-surface noise caused by bubbles and vessel-induced turbulence. Noise spikes and anomalous echoes near the seabed were removed by visual inspection. Acoustic data collected during Navigation A in May 2025 were excluded from analyses due to excessive noise caused by instrument malfunction.

To evaluate differences in overall fish biomass between the F-OWF and control areas, vertical NASC per mile values were first integrated across all depth bins within each survey, yielding a total NASC per mile for each voyage. These voyage-level biomass indices were then compared between areas. Because total NASC values satisfied the assumption of normality (Shapiro–Wilk test: W = 0.86, p = 0.06), differences between the F-OWF and control areas were assessed using a parametric t-test. The relationship between water depth and NASC per mile was further examined separately for the F-OWF and control areas to characterize vertical patterns in fish aggregation. As NASC per mile values did not meet normality assumptions (Shapiro–Wilk test: W = 0.49, p = 2.20 × 10^−16^), Spearman’s rank correlation coefficients were calculated. Because absolute water depths differed slightly between the two areas, direct comparisons based on raw depth values were not appropriate. Therefore, depth was normalized within each station by assigning a value of 1 to the sea surface and 0 to the local seabed (i.e., maximum depth). Using this normalized scale, a weighted mean normalized depth (WMND) was calculated for each survey. WMND allowed comparison of the relative vertical distribution of fish aggregations between the F-OWF and control areas while controlling for differences in absolute depth.

All statistical analyses were conducted using R version 4.4.2 (R Core Team, 2024), with statistical significance evaluated at an alpha level of 0.05.

## RESULTS

Environmental DNA metabarcoding detected at least one fish taxon in 94 out of 120 samples (78.3%). In total, 126 fish species were identified across all samples, with 95 species detected in the F-OWF area and 112 species in the control area (Supplementary Table S3). Species richness varied seasonally, with the lowest number of detected species observed in April 2024 (45 species) and the highest in August 2025 (89 species; Fig. 4a). Across depth layers, 67 species were detected in surface waters, 82 species in mid-water, and 114 species in bottom layers (Fig. 4a). Mean species richness per sample was 20.1 ± 19.5 (SD) in the F-OWF area and 25.7 ± 22.8 (SD) in the control area. However, mean species richness did not differ significantly between areas (p = 0.18; Fig. 4b). In contrast, GLM analysis revealed that water layer, sampling period, and area type all had statistically significant effects on species richness. In addition, a significant interaction between water layer and sampling period was detected (Table 1), indicating that vertical patterns in species richness varied seasonally.

**Table 1.**
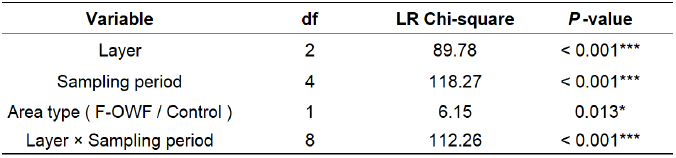
Summary of generalized linear model (GLM) results evaluating factors influencing the number of fish species detected per sample. Asterisks indicate levels of statistical significance (*p < 0.05, ***p < 0.001).

| Variable | df | LR Chi-square | P-value |
| --- | --- | --- | --- |
| Layer | 2 | 89.78 | < 0.001*** |
| Sampling period | 4 | 118.27 | < 0.001*** |
| Area type ( F-OWF / Control ) | 1 | 6.15 | 0.013* |
| Layer × Sampling period | 8 | 112.26 | < 0.001*** |

**Fig. 4.**
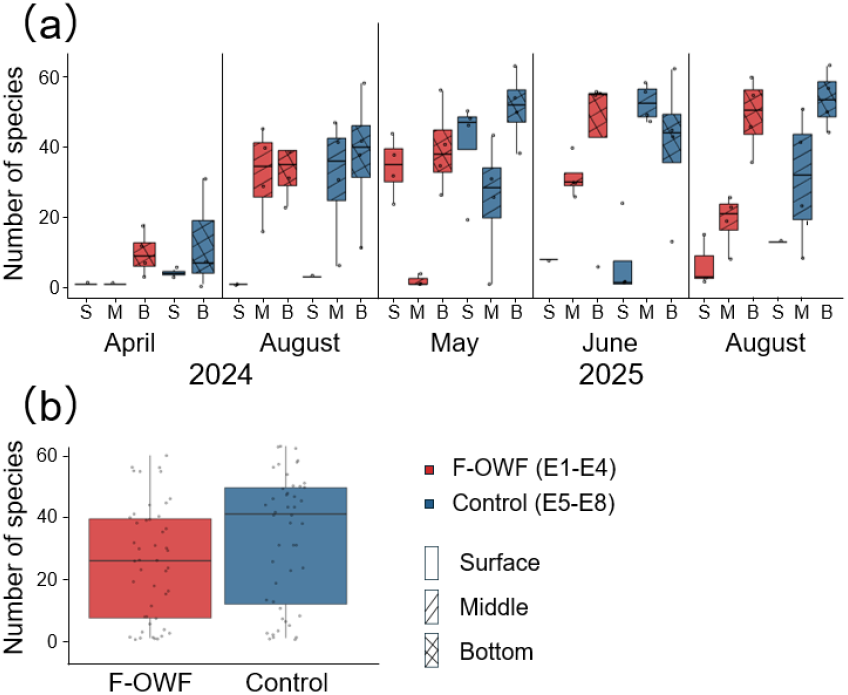
Boxplots of the number of fish species detected under different survey areas and water layers. F-OWF sites are shown in red and control sites in blue. The central line indicates the median, and the lower and upper edges of the box represent the first and third quartiles (interquartile range, IQR), respectively. (a) Seasonal and depth-specific variation in the number of detected fish species across sampling periods and water layers (S: surface, M: middle, B: bottom). Different box patterns indicate different water layers. (b) Number of fish species detected per sample in the F-OWF and control areas, with data pooled across all sampling periods and water layers.

The three most frequently detected taxa based on eDNA read counts were Dorab wolf-herring (*Chirocentrus dorab*, 1,122 reads), spotted scat (*Scatophagus argus*, 1,055 reads), and Nakahara’s longfin (*Plesiops nakaharae*, 817 reads; Fig. 5a). Four commercially important species subject to total allowable catch (TAC) management were detected: Japanese jack mackerel (*Trachurus japonicus*, 143 reads), Japanese anchovy (*Engraulis japonicus*, 2 reads), Alaska pollock (*Gadus chalcogrammus*, 59 reads), and blue mackerel (*Scomber australasicus*, 188 reads). Stacked bar charts illustrating relative species dominance showed broadly similar community composition between the F-OWF and control areas throughout the year (Fig. 5a). This pattern remained consistent when data were pooled across all sampling periods (Fig. 5b). Consistent with these observations, PERMANOVA based on Bray–Curtis dissimilarity indicated no significant difference in overall fish community structure between the F-OWF and control areas (R^2^ = 0.13, p = 0.60; Table 2).

**Table 2.** PERMANOVA results based on Bray–Curtis dissimilarity for fish community structure between wind turbine and control stations. Fish community composition was analyzed using PERMANOVA based on Bray–Curtis dissimilarity. Prior to analysis, read counts were normalized so that the cumulative read abundance of the 30 most abundant taxa across all stations summed to one, representing relative dominance within the top 30 taxa. The PERMANOVA results indicated no significant difference in fish community structure between wind turbine and control stations.

| Source | df | R<sup>2</sup> | F | P-value |
| --- | --- | --- | --- | --- |
| Area type ( F-OWF / Control ) | 1 | 0.13 | 0.92 | 0.60 |
| Residuals | 6 | 0.87 |  |  |
| Total | 7 | 1 |  |  |

**Fig. 5.**
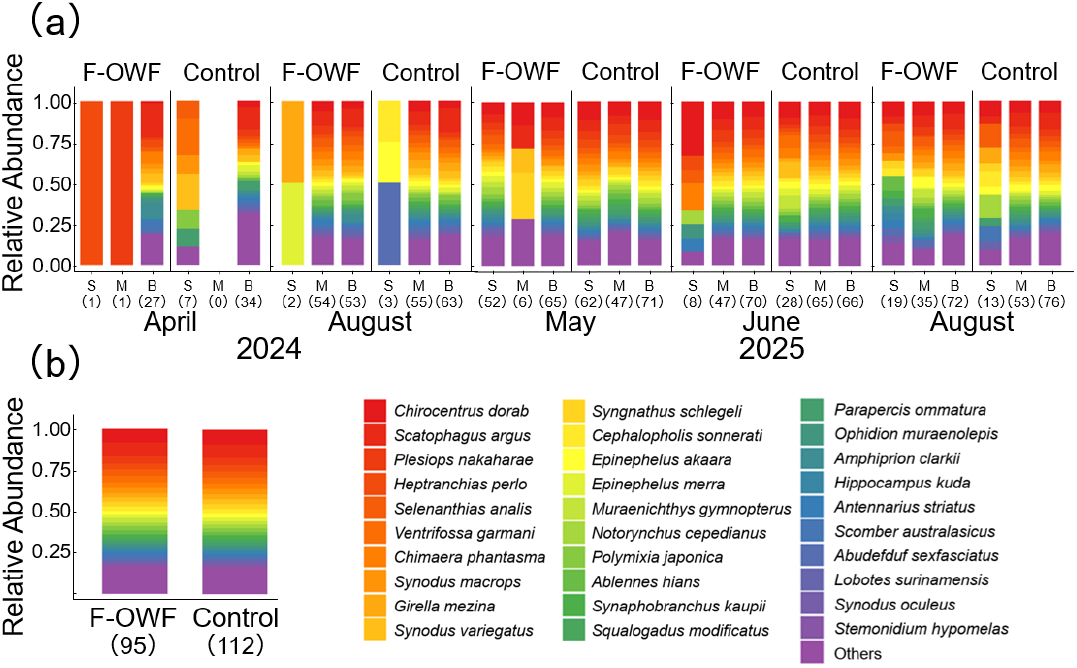
Heatmap showing the relative dominance of the 30 most frequently detected fish species based on eDNA metabarcoding. Values in parentheses indicate the number of detected fish species. (a) Relative dominance of fish species across survey areas (F-OWF and control), water layers (S: surface, M: middle, B: bottom), and sampling periods (April and August 2024; May, June, and August 2025). (b) Overall relative dominance of detected fish species in the F-OWF and control areas, with data pooled across all water layers and sampling periods, illustrating broadly similar community composition between areas.

Representative echograms obtained from the scientific echosounder surveys are shown in Fig. 6. Across all surveys, NASC per mile values ranged from a minimum of 0 to a maximum of 461.5, with a mean value of 26.7 (Fig. 7a). Spearman’s rank correlation analysis revealed a significant relationship between water depth and NASC per mile in the control area (p = 0.001), whereas no such relationship was detected in the F-OWF area (p = 0.80). These results indicate a fundamental difference in the vertical structuring of fish aggregations between areas with and without floating offshore wind turbines. When vertically integrated NASC per mile values were compared at the voyage level, mean total NASC per mile (± SD) was 352.6 ± 295.1 in the control area and 393.1 ± 304.7 in the F-OWF area (Fig. 7b). Although this difference was not statistically significant (p = 0.83), total NASC per mile tended to be 11.5% higher in the F-OWF area (Fig. 7b). The weighted mean normalized depth (WMND) of fish aggregations differed between areas: WMND was 0.34 in the F-OWF area (corresponding to a mean depth of 87.9 m) and 0.24 in the control area (corresponding to a mean depth of 139 m) (Fig. 7c). This indicates that the vertical center of fish aggregation in the F-OWF area was approximately 13.2% shallower relative to the water column than in the control area. All echograms and detailed NASC metrics are provided in Supplementary Fig. S1 and Supplementary Table S4.

**Fig. 6.**
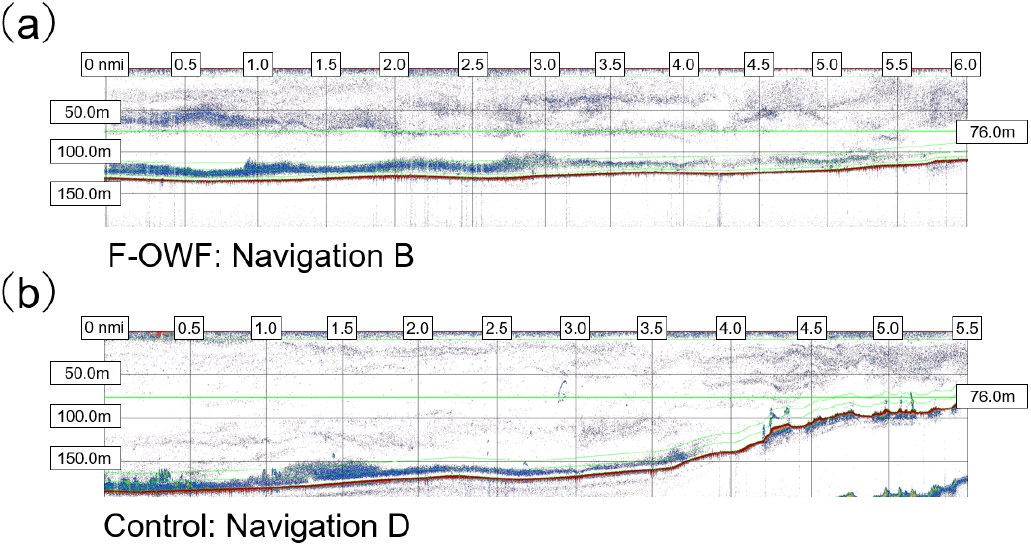
Representative echograms obtained using a scientific echosounder. (a) Echogram from the wind turbine station during transect B in May. (b) Echogram from the control station during transect D in May. Acoustic data were collected using a scientific echosounder (EK80; Simrad Kongsberg Maritime AS, Horten, Norway) operated in continuous-wave (CW) mode with a transmit power of 150 W, frequency of 200 kHz, pulse duration of 1.024 ms, beam width of 7°, and ping interval of 1000 ms. Echograms were visualized using Echoview software (version 9.0; Echoview Software Pty Ltd). Colors represent volume backscattering strength (Sv) displayed using the standard color scale. The horizontal dark green band observed in the mid-water layer indicates the draft depth of the wind turbine structure (76 m).

**Fig. 7.**
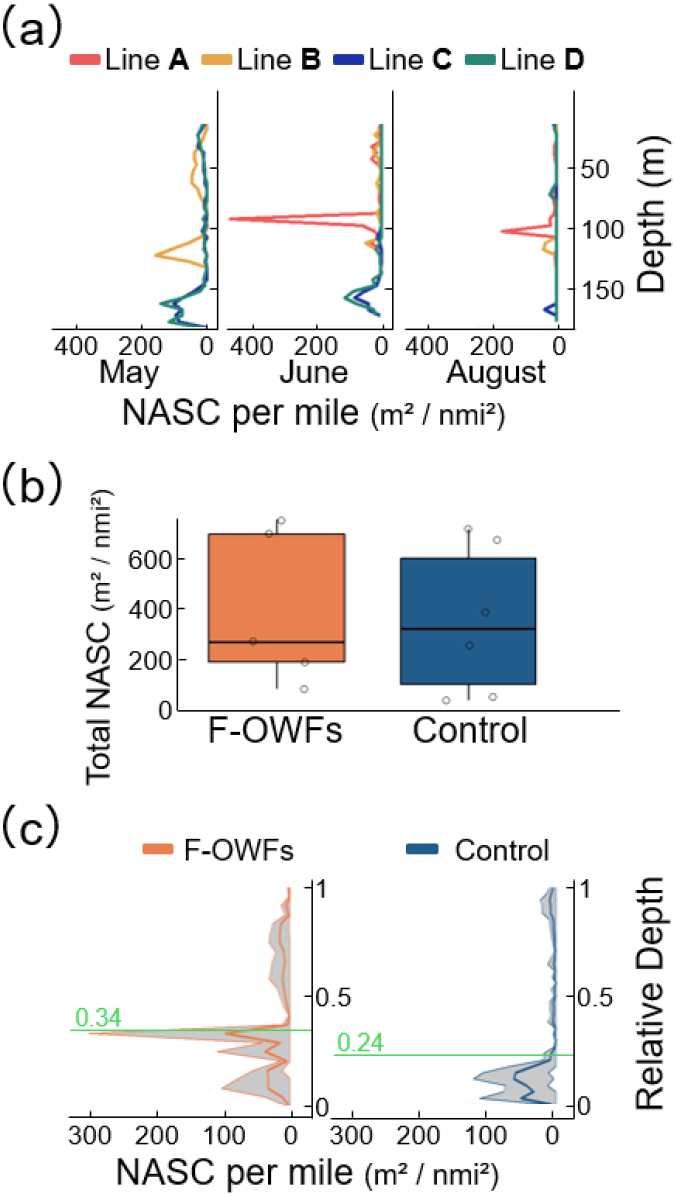
Fish density patterns in the floating offshore wind farm (F-OWF) and control areas based on acoustic measurements. (a) Vertical profiles of NASC per mile for each transect and sampling month. In the F-OWF area, Line A is shown in red and Line B in ochre; in the control area, Line C is shown in blue and Line D in green. Data from Line A in May were excluded due to noise caused by an acoustic instrument malfunction. (b) Total NASC per mile integrated over the water column for each transect, compared between the F-OWF (orange) and control (blue) areas. Box plots show the median (center line), first and third quartiles (box limits), and interquartile range (whiskers); circles represent individual transect data points. (c) Mean vertical distribution of NASC per mile across all sampling periods for the F-OWF (orange) and control (blue) areas. The y-axis represents relative water depth. Solid lines indicate mean values at each depth, and shaded areas denote standard errors (SE). The green horizontal line and accompanying value indicate the weighted mean normalized depth (WMND) for each area.

## DISCUSSION

Floating offshore wind farms (F-OWFs) are rapidly expanding into offshore pelagic environments, yet their ecological effects remain poorly constrained due to the limited number of operational sites and the logistical difficulty of conducting non-invasive monitoring at sea (Farr et al., 2021; Rezaei et al., 2023). Here, we combined eDNA metabarcoding and scientific echosounding to provide a multidimensional assessment of fish communities around the Goto Islands F-OWF. This integrated framework leverages the complementary strengths of the two approaches—taxonomic resolution from eDNA and quantitative information on aggregation structure from acoustics—and is particularly well suited for evaluating offshore infrastructures where conventional net-based sampling is constrained (Simmonds and MacLennan, 2008; Miya, 2022). Our results revealed a clear contrast: fish community composition inferred from eDNA was broadly similar between the F-OWF and control areas, whereas acoustic data indicated a consistent shift in the vertical structuring of fish aggregations in the vicinity of floating turbines.

Across sampling periods and depth layers, eDNA metabarcoding detected 126 fish species, and multivariate community structure based on the most frequently detected taxa did not differ significantly between the F-OWF and control areas. Similar findings have been reported for bottom-fixed OWFs, where dominant fish assemblages often remain comparable to surrounding reference areas despite local aggregation effects (Wilhelmsson et al., 2006; Degraer et al., 2020). The absence of a detectable compositional shift therefore does not imply the absence of ecological effects; rather, it suggests that turbine-associated influences may operate primarily through redistribution of individuals rather than replacement of dominant taxa.

Species richness varied strongly with sampling period and water layer, and the significant interaction between water layer and sampling period indicates that vertical richness patterns were seasonally dynamic. Such patterns are consistent with known constraints on eDNA detectability in offshore surface waters, where elevated temperature and ultraviolet radiation accelerate DNA degradation (Strickler et al., 2015; Lamb et al., 2022), and hydrodynamic processes promote dilution and transport away from the source (Andruszkiewicz et al., 2019). The lower detection frequency observed in surface samples, together with higher richness in bottom layers, underscores the importance of explicitly accounting for depth-related and hydrographic processes when interpreting eDNA-based spatial comparisons in open marine systems (Maruyama et al., 2014).

In contrast to the largely stable eDNA-inferred composition, acoustic surveys revealed pronounced differences in vertical distribution patterns. In the control area, fish biomass increased with depth, a pattern consistent with observations from structurally simple offshore environments where fish tend to concentrate near the seabed (Auster et al., 1991; Sutton et al., 2008). Near the F-OWFs, however, this depth-dependence disappeared, and the weighted mean normalized depth (WMND) indicated a shallower vertical center of aggregation relative to the water column.

Because WMND normalizes depth within each station, it enables comparison of relative vertical positioning while controlling for differences in absolute depth between areas (Dornan et al., 2019). The observed shallowing of aggregation depth near the turbines suggests that F-OWFs are associated with a reorganization of fish distribution in three-dimensional space, even in the absence of major changes in species composition.

A plausible mechanism is that floating turbines and their mooring systems introduce persistent physical structure and shading into the upper water column, providing orientation cues, shelter, or meeting points for pelagic fishes. Similar mechanisms have long been proposed for fish aggregating devices (FADs), where attraction can occur through non-trophic pathways such as refuge provision and facilitation of schooling behavior (Castro et al., 2002; Fréon and Dagorn, 2000). Pelagic species detected in this study, including Trachurus japonicus and Scomber australasicus, are known to associate with floating structures (Okamoto, 1992; Tsuchida et al., 2025). Conceptual syntheses of OWF effects have likewise suggested that pelagic fishes may be attracted to offshore structures primarily for non-feeding-related reasons (Degraer et al., 2020). Because our study is based on spatial comparisons rather than a before–after design, we interpret these patterns as associations with F-OWF presence rather than definitive evidence of causation.

The decoupling between taxonomic composition and vertical structure likely reflects the high mobility of pelagic fishes and the spatial scale of observation. If floating structures primarily concentrate individuals from the surrounding seascape without excluding taxa, community composition may remain broadly similar while aggregation intensity and vertical positioning shift (Wilhelmsson et al., 2006; Degraer et al., 2020). This interpretation is consistent with the non-significant difference in voyage-level integrated NASC per mile between areas, despite a higher mean value near turbines, indicating redistribution rather than a net increase in biomass within the surveyed footprint. Several limitations warrant consideration. First, eDNA metabarcoding provides robust information on species presence but has inherent constraints for quantitative inference due to primer bias, amplification stochasticity, and environmental variability in DNA transport and degradation (Kebschull and Zador, 2015; Collins et al., 2019). Second, acoustic estimates may underestimate demersal fish biomass due to the acoustic dead zone near the seabed (Ona and Mitson, 1996), highlighting the value of complementary visual validation using remotely operated vehicles or optical systems. Third, ecological responses to offshore structures can evolve over multi-year timescales as communities undergo successional changes (Stenberg et al., 2015; Degraer et al., 2020). Long-term post-construction monitoring will therefore be essential to determine whether the observed redistribution persists, intensifies, or ultimately translates into changes in community composition.

## Conclusions

By integrating eDNA metabarcoding and scientific echosounding, this study demonstrates that floating offshore wind infrastructure can be associated with changes in the three-dimensional spatial organization of fish aggregations, even when dominant taxonomic composition remains broadly similar to nearby control areas. These findings highlight the importance of coupling taxonomic and spatial metrics when evaluating offshore renewable energy infrastructures and underscore the need for standardized, long-term monitoring to support adaptive management as F-OWF deployment accelerates worldwide.

## Acknowledgments

We sincerely thank the crew of the training vessel (T/S) Kakuyo-Maru for their great support in this research. We also appreciate the staff of the Fish and Ships Laboratory, Faculty of Fisheries, Nagasaki University.

## Ethics Statement

The research required no permit approvals.

## Data Availability Statement

The data that supports the findings of this study are available in the supplementary material of this article.

