## Supplementary Information for "Three-dimensional redistribution of pelagic fish aggregations associated with floating offshore wind farms"

Table S1 Geographic coordinates and water depths of sampling stations and wind turbines. Latitude and longitude of each sampling station and wind turbine are provided together with water depth. For wind turbines, the construction year is also included.

| F-OWFs/Station | Latitude | Longitude | Depth (m) | Construction Year |
|---|---|---|---|---|
| **FH** | 32º 40.279'N | 128º 56.32'E | 100 | 2015 |
| **F1** | 32º 40.333'N | 128º 57.455'E | 120 | 2022 |
| **F2** | 32º 40.332'N | 128º 57.845'E | 120 | 2022 |
| **F3** | 32º 40.335'N | 128º 58.237'E | 125 | 2022 |
| **F4** | 32º 40.192'N | 128º 58.637'E | 130 | 2025 |
| **F5** | 32º 40.232'N | 128º 59.089'E | 130 | 2025 |
| **F6** | 32º 40.138'N | 128º 59.451'E | 135 | 2025 |
| **F7** | 32º 40.130'N | 128º 59.841'E | 135 | 2025 |
| **F8** | 32º 40.276'N | 129º 00.166'E | 135 | 2025 |
| **E1** | 32º 40.5'N | 129º 0.3'E | 119 | - |
| **E2** | 32º 40.5'N | 128º 59.3'E | 128 | - |
| **E3** | 32º 40.5'N | 128º 58.3'E | 131 | - |
| **E4** | 32º 40.5'N | 128º 57.3'E | 133 | - |
| **E5** | 32º 36.5'N | 128º 57.3'E | 168 | - |
| **E6** | 32º 36.5'N | 128º 58.3'E | 160 | - |
| **E7** | 32º 36.5'N | 128º 59.3'E | 140 | - |
| **E8** | 32º 36.5'N | 129º 0.3'E | 87 | - |
| **S1** | 32º 36.0'N | 128º 57.290'E | 85 | - |
| **S2** | 32º 36.0'N | 128º 57.890'E | 115 | - |
| **S3** | 32º 36.0'N | 128º 58.490'E | 140 | - |
| **S4** | 32º 36.0'N | 128º 59.090'E | 160 | - |
| **S5** | 32º 36.0'N | 128º 59.690'E | 165 | - |
| **S6** | 32º 36.0'N | 129º 00.290'E | 170 | - |

Table S2 CTD採水機により取得した環境要因。サンプリング時間、採水場所、水深（m）、水温（℃）、塩分を示す。

| Sampling Time | Area | Site | Layer | Water Depth (m) | Temperature (℃) | Salinity |
|---|---|---|---|---|---|---|
| 2024/4/25 8:24 | F-OWF | E1 | Surface | 5 | 18.5144 | 34.4651 |
| 2024/4/25 8:24 | F-OWF | E1 | Middle | 50 | 17.3979 | 34.4363 |
| 2024/4/25 8:24 | F-OWF | E1 | Bottom | 115 | 15.0208 | 34.5149 |
| 2024/4/25 7:59 | F-OWF | E2 | Surface | 5 | 17.9675 | 34.4387 |
| 2024/4/25 7:59 | F-OWF | E2 | Middle | 50 | 17.4441 | 34.4425 |
| 2024/4/25 7:59 | F-OWF | E2 | Bottom | 123 | 14.5685 | 34.503 |
| 2024/4/25 7:43 | F-OWF | E3 | Surface | 5 | 18.383 | 34.4519 |
| 2024/4/25 7:43 | F-OWF | E3 | Middle | 50 | 17.4041 | 34.4319 |
| 2024/4/25 7:43 | F-OWF | E3 | Bottom | 125 | 15.0702 | 34.5171 |
| 2024/4/25 7:24 | F-OWF | E4 | Surface | 5 | 17.7851 | 34.2813 |
| 2024/4/25 7:24 | F-OWF | E4 | Middle | 50 | 17.1226 | 34.3768 |
| 2024/4/25 7:24 | F-OWF | E4 | Bottom | 125 | 14.7279 | 34.5074 |
| 2024/4/25 9:29 | Control | E5 | Surface | 5 | 18.6379 | 34.4863 |
| 2024/4/25 9:29 | Control | E5 | Middle | 50 | 17.6813 | 34.5317 |
| 2024/4/25 9:29 | Control | E5 | Bottom | 165 | 12.8086 | 34.4438 |
| 2024/4/25 9:14 | Control | E6 | Surface | 5 | 18.6456 | 34.4862 |
| 2024/4/25 9:14 | Control | E6 | Middle | 50 | 17.5266 | 34.5505 |
| 2024/4/25 9:14 | Control | E6 | Bottom | 155 | 13.6081 | 34.4719 |
| 2024/4/25 8:59 | Control | E7 | Surface | 5 | 18.6517 | 34.4831 |
| 2024/4/25 8:59 | Control | E7 | Middle | 50 | 17.6843 | 34.5337 |
| 2024/4/25 8:59 | Control | E7 | Bottom | 82 | 14.1665 | 34.4912 |

| | | | | | | |
|---|---|---|---|---|---|---|
| 2024/4/25 8:46 | Control | E8 | Surface | 5 | 18.6493 | 34.4817 |
| 2024/4/25 8:46 | Control | E8 | Middle | 50 | 17.4778 | 34.4418 |
| 2024/4/25 8:46 | Control | E8 | Bottom | 82 | 16.7092 | 34.4836 |
| 2024/8/15 8:33 | F-OWF | E1 | Surface | 5 | 29.9292 | 31.3022 |
| 2024/8/15 8:33 | F-OWF | E1 | Middle | 50 | 19.2737 | 34.0629 |
| 2024/8/15 8:33 | F-OWF | E1 | Bottom | 113 | 16.6415 | 34.4937 |
| 2024/8/15 8:03 | F-OWF | E2 | Surface | 5 | 30.3183 | 31.1334 |
| 2024/8/15 8:03 | F-OWF | E2 | Middle | 50 | 18.9639 | 34.1685 |
| 2024/8/15 8:03 | F-OWF | E2 | Bottom | 121 | 16.5179 | 34.5079 |
| 2024/8/15 7:46 | F-OWF | E3 | Surface | 5 | 29.9826 | 31.4733 |
| 2024/8/15 7:46 | F-OWF | E3 | Middle | 50 | 20.1331 | 33.9252 |
| 2024/8/15 7:46 | F-OWF | E3 | Bottom | 125 | 16.22 | 34.5078 |
| 2024/8/15 7:27 | F-OWF | E4 | Surface | 5 | 30.4488 | 31.0535 |
| 2024/8/15 7:27 | F-OWF | E4 | Middle | 50 | 19.0213 | 34.1371 |
| 2024/8/15 7:27 | F-OWF | E4 | Bottom | 127 | 15.9531 | 34.5043 |
| 2024/8/15 9:56 | Control | E5 | Surface | 5 | 30.815 | 30.5795 |
| 2024/8/15 9:56 | Control | E5 | Middle | 50 | 19.8658 | 33.9255 |
| 2024/8/15 9:56 | Control | E5 | Bottom | 163 | 14.205 | 34.4767 |
| 2024/8/15 9:30 | Control | E6 | Surface | 5 | 30.7827 | 30.7374 |
| 2024/8/15 9:30 | Control | E6 | Middle | 50 | 20.5137 | 33.7741 |
| 2024/8/15 9:30 | Control | E6 | Bottom | 155 | 14.9243 | 34.4968 |
| 2024/8/15 9:15 | Control | E7 | Surface | 5 | 30.8377 | 30.7009 |
| 2024/8/15 9:15 | Control | E7 | Middle | 50 | 20.6761 | 33.7204 |

| | | | | | | |
|---|---|---|---|---|---|---|
| 2024/8/15 9:15 | Control | E7 | Bottom | 135 | 16.168 | 34.5038 |
| 2024/8/15 9:01 | Control | E8 | Surface | 5 | 30.3592 | 31.1746 |
| 2024/8/15 9:01 | Control | E8 | Middle | 50 | 20.3126 | 33.8055 |
| 2024/8/15 9:01 | Control | E8 | Bottom | 81 | 16.7707 | 34.4585 |
| 2025/5/1 13:11 | F-OWF | E1 | Surface | 5 | 17.0495 | 34.5954 |
| 2025/5/1 13:11 | F-OWF | E1 | Middle | 50 | 16.1163 | 34.5799 |
| 2025/5/1 13:11 | F-OWF | E1 | Bottom | 110 | 14.4236 | 34.5364 |
| 2025/5/1 13:30 | F-OWF | E2 | Surface | 5 | 17.8861 | 34.6126 |
| 2025/5/1 13:30 | F-OWF | E2 | Middle | 50 | 16.3602 | 34.5867 |
| 2025/5/1 13:30 | F-OWF | E2 | Bottom | 120 | 14.3163 | 34.5326 |
| 2025/5/1 13:50 | F-OWF | E3 | Surface | 5 | 17.8104 | 34.617 |
| 2025/5/1 13:50 | F-OWF | E3 | Middle | 50 | 16.3504 | 34.5855 |
| 2025/5/1 13:50 | F-OWF | E3 | Bottom | 125 | 14.3157 | 34.5322 |
| 2025/5/1 14:11 | F-OWF | E4 | Surface | 5 | 18.0237 | 34.6157 |
| 2025/5/1 14:11 | F-OWF | E4 | Middle | 50 | 16.4174 | 34.5849 |
| 2025/5/1 14:11 | F-OWF | E4 | Bottom | 125 | 14.4639 | 34.5395 |
| 2025/5/1 10:02 | Control | E5 | Surface | 5 | - | - |
| 2025/5/1 10:02 | Control | E5 | Middle | 50 | - | - |
| 2025/5/1 10:02 | Control | E5 | Bottom | 160 | - | - |
| 2025/5/1 10:24 | Control | E6 | Surface | 5 | - | - |
| 2025/5/1 10:24 | Control | E6 | Middle | 50 | - | - |
| 2025/5/1 10:24 | Control | E6 | Bottom | 157 | - | - |
| 2025/5/1 10:44 | Control | E7 | Surface | 5 | 17.7576 | 34.6051 |

| | | | | | | |
|---|---|---|---|---|---|---|
| 2025/5/1 10:44 | Control | E7 | Middle | 50 | 16.4294 | 34.5876 |
| 2025/5/1 10:44 | Control | E7 | Bottom | 140 | 14.5175 | 34.5428 |
| 2025/5/1 11:05 | Control | E8 | Surface | 5 | 17.567 | 34.6086 |
| 2025/5/1 11:05 | Control | E8 | Middle | 50 | 16.4041 | 34.586 |
| 2025/5/1 11:05 | Control | E8 | Bottom | 85 | 14.6539 | 34.5397 |
| 2025/6/5 7:16 | F-OWF | E1 | Surface | 5 | 19.339 | 34.1415 |
| 2025/6/5 7:16 | F-OWF | E1 | Middle | 50 | 18.6965 | 34.3062 |
| 2025/6/5 7:16 | F-OWF | E1 | Bottom | 110 | 16.6582 | 34.5592 |
| 2025/6/5 6:59 | F-OWF | E2 | Surface | 5 | 19.8626 | 33.9764 |
| 2025/6/5 6:59 | F-OWF | E2 | Middle | 50 | 18.2556 | 34.4541 |
| 2025/6/5 6:59 | F-OWF | E2 | Bottom | 120 | 16.5712 | 34.5648 |
| 2025/6/5 6:42 | F-OWF | E3 | Surface | 5 | 20.1899 | 33.6977 |
| 2025/6/5 6:42 | F-OWF | E3 | Middle | 50 | 17.967 | 34.4801 |
| 2025/6/5 6:42 | F-OWF | E3 | Bottom | 125 | 16.3957 | 34.5684 |
| 2025/6/5 6:24 | F-OWF | E4 | Surface | 5 | 20.2109 | 33.6818 |
| 2025/6/5 6:24 | F-OWF | E4 | Middle | 50 | 17.7032 | 34.5097 |
| 2025/6/5 6:24 | F-OWF | E4 | Bottom | 125 | 16.4005 | 34.5689 |
| 2025/6/5 10:02 | Control | E5 | Surface | 5 | 20.2749 | 33.8038 |
| 2025/6/5 10:02 | Control | E5 | Middle | 50 | 16.9351 | 34.473 |
| 2025/6/5 10:02 | Control | E5 | Bottom | 165 | 15.0772 | 34.5538 |
| 2025/6/5 9:45 | Control | E6 | Surface | 5 | 20.2244 | 33.7608 |
| 2025/6/5 9:45 | Control | E6 | Middle | 50 | 17.0266 | 34.5381 |
| 2025/6/5 9:45 | Control | E6 | Bottom | 155 | 15.6218 | 34.5644 |

| | | | | | | |
|---|---|---|---|---|---|---|
| 2025/6/5 9:29 | Control | E7 | Surface | 5 | 20.2355 | 33.9036 |
| 2025/6/5 9:29 | Control | E7 | Middle | 50 | 17.2422 | 34.5317 |
| 2025/6/5 9:29 | Control | E7 | Bottom | 133 | 16.2473 | 34.5701 |
| 2025/6/5 9:14 | Control | E8 | Surface | 5 | 19.974 | 33.9569 |
| 2025/6/5 9:14 | Control | E8 | Middle | 50 | 17.6329 | 34.5062 |
| 2025/6/5 9:14 | Control | E8 | Bottom | 83 | 16.6576 | 34.563 |
| 2025/8/20 15:42 | F-OWF | E1 | Surface | 10 | 27.5517 | 33.5949 |
| 2025/8/20 15:42 | F-OWF | E1 | Middle | 50 | 25.3368 | 33.8005 |
| 2025/8/20 15:42 | F-OWF | E1 | Bottom | 105 | 17.5383 | 34.5365 |
| 2025/8/20 15:59 | F-OWF | E2 | Surface | 10 | 27.5661 | 33.6129 |
| 2025/8/20 15:59 | F-OWF | E2 | Middle | 50 | 25.0567 | 33.8148 |
| 2025/8/20 15:59 | F-OWF | E2 | Bottom | 115 | 16.9421 | 34.547 |
| 2025/8/20 16:17 | F-OWF | E3 | Surface | 10 | 28.1149 | 33.6617 |
| 2025/8/20 16:17 | F-OWF | E3 | Middle | 50 | 25.8615 | 33.7369 |
| 2025/8/20 16:17 | F-OWF | E3 | Bottom | 120 | 17.1858 | 34.5453 |
| 2025/8/20 16:34 | F-OWF | E4 | Surface | 10 | 28.3487 | 33.6501 |
| 2025/8/20 16:34 | F-OWF | E4 | Middle | 50 | 24.8972 | 33.8385 |
| 2025/8/20 16:34 | F-OWF | E4 | Bottom | 120 | 17.2829 | 34.5458 |
| 2025/8/20 12:46 | Control | E5 | Surface | 5 | 28.5803 | 33.6448 |
| 2025/8/20 12:46 | Control | E5 | Middle | 50 | 25.7332 | 33.7357 |
| 2025/8/20 12:46 | Control | E5 | Bottom | 165 | 14.9953 | 34.5177 |
| 2025/8/20 13:10 | Control | E6 | Surface | 5 | 28.5538 | 33.62 |
| 2025/8/20 13:10 | Control | E6 | Middle | 50 | 25.6232 | 33.7626 |

| | | | | | | |
|---|---|---|---|---|---|---|
| 2025/8/20 13:10 | Control | E6 | Bottom | 150 | 16.0328 | 34.5397 |
| 2025/8/20 13:33 | Control | E7 | Surface | 5 | 28.0055 | 33.6395 |
| 2025/8/20 13:33 | Control | E7 | Middle | 50 | 25.8271 | 33.7441 |
| 2025/8/20 13:33 | Control | E7 | Bottom | 130 | 16.6526 | 34.5421 |
| 2025/8/20 13:52 | Control | E8 | Surface | 10 | 28.2916 | 33.5817 |
| 2025/8/20 13:52 | Control | E8 | Middle | 50 | 25.7546 | 33.7404 |
| 2025/8/20 13:52 | Control | E8 | Bottom | 75 | 21.326 | 34.2622 |
| 2024/4/25 9:40 | - | BLANK | - | - | - | - |
| 2024/8/15 9:56 | - | BLANK | - | - | - | - |
| 2025/5/1 14:11 | - | BLANK | - | - | - | - |
| 2025/6/5 10:02 | - | BLANK | - | - | - | - |
| 2025/8/20 16:34 | - | BLANK | - | - | - | - |
| - | - | 1st_BLANK | - | - | - | - |

Table S3 Fish species detected by eDNA metabarcoding and their detection frequencies.

| **Area** | **Rank** | **Scientific name** | **Common name** | **Total Frequency** |
|---|---|---|---|---|
| F-OWF | 1 | *Scatophagus argus* | Spotted scat | 403 |
| F-OWF | 2 | *Chirocentrus dorab* | Dorab wolf-herring | 394 |
| F-OWF | 3 | *Plesiops nakaharae* | Nakahara's longfin | 209 |
| F-OWF | 4 | *Ventrifossa garmani* | Sagami grenadier | 194 |
| F-OWF | 5 | *Selenanthias analis* | Pearl-spotted fairy basslet | 172 |
| F-OWF | 6 | *Heptranchias perlo* | Sharpnose sevengill shark | 170 |
| F-OWF | 7 | *Chimaera phantasma* | Silver chimaera | 158 |
| F-OWF | 8 | *Synodus macrops* | Triplecross lizardfish | 155 |

| | | | | |
|---|---|---|---|---|
| F-OWF | 9 | *Muraenichthys gymnopterus* | Anguilliformes Ophichthidae | 134 |
| F-OWF | 10 | *Synodus variegatus* | Variegated lizardfish | 131 |
| F-OWF | 11 | *Syngnathus schlegeli* | Seaweed pipefish | 130 |
| F-OWF | 12 | *Girella mezina* | Yellowstriped blackfish | 125 |
| F-OWF | 13 | *Cephalopholis sonnerati* | Tomato hind | 121 |
| F-OWF | 14 | *Polymixia japonica* | Silver eye | 114 |
| F-OWF | 15 | *Ablennes hians* | Flat needlefish | 111 |
| F-OWF | 16 | *Epinephelus merra* | Honeycomb grouper | 111 |
| F-OWF | 17 | *Notorynchus cepedianus* | Broadnose sevengill shark | 105 |
| F-OWF | 18 | *Epinephelus akaara* | Hong Kong grouper | 98 |
| F-OWF | 19 | *Synaphobranchus kaupii* | Kaup's arrowtooth eel | 95 |
| F-OWF | 20 | *Amphiprion clarkii* | Yellowtail clownfish | 93 |
| F-OWF | 21 | *Parapercis ommatura* | Perciformes: Pinguipedidae | 88 |
| F-OWF | 22 | *Abudefduf sexfasciatus* | Scissortail sergeant | 85 |
| F-OWF | 23 | *Squalogadus modificatus* | Tadpole whiptail | 84 |
| F-OWF | 24 | *Antennarius striatus* | Striated frogfish | 82 |
| F-OWF | 25 | *Ophidion muraenolepis* | Blackedge cusk | 75 |
| F-OWF | 26 | *Scomber australasicus* | Blue mackerel | 75 |
| F-OWF | 27 | *Hippocampus kuda* | Spotted seahorse | 71 |
| F-OWF | 28 | *Eurypharynx pelecanoides* | Pelican eel | 69 |
| F-OWF | 29 | *Stemonidium hypomelas* | Black serrivomerid eel | 59 |
| F-OWF | 30 | *Trachurus japonicus* | Japanese jack mackerel | 59 |
| F-OWF | 31 | *Lobotes surinamensis* | Tripletail | 58 |
| F-OWF | 32 | *Synodus oculeus* | Large-eye lizardfish | 56 |

| | | | | |
|---|---|---|---|---|
| F-OWF | 33 | *Corythoichthys flavofasciatus* | Network pipefish | 48 |
| F-OWF | 34 | *Oxyconger leptognathus* | Shorttail pike conger | 47 |
| F-OWF | 35 | *Amblyglyphidodon curacao* | Staghorn damselfish | 46 |
| F-OWF | 36 | *Gymnothorax kidako* | Kidako moray | 46 |
| F-OWF | 37 | *Harriotta raleighana* | Pacific longnose chimaera | 43 |
| F-OWF | 38 | *Sigmops gracilis* | Slender fangjaw | 39 |
| F-OWF | 39 | *Sphyraena pinguis* | Red barracuda | 34 |
| F-OWF | 40 | *Herklotsichthys quadrimaculatus* | Bluestripe herring | 33 |
| F-OWF | 41 | *Valenciennellus tripunctulatus* | Constellationfish | 33 |
| F-OWF | 42 | *Plectroglyphidodon leucozonus* | Singlebar devil | 28 |
| F-OWF | 43 | *Euthynnus affinis* | Kawakawa | 27 |
| F-OWF | 44 | *Cynoglossus bilineatus* | Fourlined tonguesole | 26 |
| F-OWF | 45 | *Albula glossodonta* | Roundjaw bonefish | 24 |
| F-OWF | 46 | *Fowlerichthys scriptissimus* | Fishing frog | 24 |
| F-OWF | 47 | *Cynoglossus joyneri* | Red tonguesole | 21 |
| F-OWF | 48 | *Gadus chalcogrammus* | Alaska pollock | 20 |
| F-OWF | 49 | *Pomacentrus brachialis* | Charcoal damsel | 16 |
| F-OWF | 50 | *Pempheris schwenkii* | Silver sweeper | 12 |
| F-OWF | 51 | *Pomacentrus moluccensis* | Lemon damsel | 8 |
| F-OWF | 52 | *Lophotus capellei* | Unicornfish | 7 |
| F-OWF | 53 | *Bolbometopon muricatum* | Green humphead parrotfish | 6 |
| F-OWF | 54 | *Girella punctata* | Largescale blackfish | 5 |
| F-OWF | 55 | *Lobianchia gemellarii* | Cocco's lantern fish | 5 |
| F-OWF | 56 | *Paraplagusia japonica* | Black cow-tongue | 5 |

| | | | | |
|---|---|---|---|---|
| F-OWF | 57 | *Vinciguerria nimbaria* | Oceanic lightfish | 5 |
| F-OWF | 58 | *Anguilla marmorata* | Giant mottled eel | 4 |
| F-OWF | 59 | *Anampses geographicus* | Geographic wrasse | 3 |
| F-OWF | 60 | *Pristipomoides filamentosus* | Crimson jobfish | 3 |
| F-OWF | 61 | *Pterygotrigla ryukyuensis* | Ryukyu gurnard | 3 |
| F-OWF | 62 | *Thunnus obesus* | Bigeye tuna | 3 |
| F-OWF | 63 | *Alepisaurus ferox* | Long snouted lancetfish | 2 |
| F-OWF | 64 | *Anyperodon leucogrammicus* | Slender grouper | 2 |
| F-OWF | 65 | *Arnoglossus polyspilus* | Many-spotted lefteye flounder | 2 |
| F-OWF | 66 | *Chlamydoselachus anguineus* | Frilled shark | 2 |
| F-OWF | 67 | *Gazza minuta* | Toothpony | 2 |
| F-OWF | 68 | *Kyphosus bigibbus* | Brown chub | 2 |
| F-OWF | 69 | *Melanocetus johnsonii* | Humpback anglerfish | 2 |
| F-OWF | 70 | *Metavelifer multiradiatus* | Spinyfin velifer | 2 |
| F-OWF | 71 | *Moringua microchir* | Lesser thrush eel | 2 |
| F-OWF | 72 | *Pomacentrus coelestis* | Neon damselfish | 2 |
| F-OWF | 73 | *Ptereleotris heteroptera* | Blacktail goby | 2 |
| F-OWF | 74 | *Zeus faber* | John dory | 2 |
| F-OWF | 75 | *Argentina kagoshimae* | - | 1 |
| F-OWF | 76 | *Atherion elymus* | Bearded silverside | 1 |
| F-OWF | 77 | *Champsodon snyderi* | Benttooth | 1 |
| F-OWF | 78 | *Cottus kazika* | - | 1 |
| F-OWF | 79 | *Engraulis japonicus* | Japanese anchovy | 1 |
| F-OWF | 80 | *Eviota afelei* | Afele's dwarfgoby | 1 |

| | | | | |
|---|---|---|---|---|
| F-OWF | 81 | *Iracundus signifer* | Decoy scorpionfish | 1 |
| F-OWF | 82 | *Johnius belangerii* | Belanger's croaker | 1 |
| F-OWF | 83 | *Lutjanus argentimaculatus* | Mangrove red snapper | 1 |
| F-OWF | 84 | *Macropharyngodon meleagris* | Blackspotted wrasse | 1 |
| F-OWF | 85 | *Nealotus tripes* | Black snake mackerel | 1 |
| F-OWF | 86 | *Neoclinus chihiroe* | - | 1 |
| F-OWF | 87 | *Paraplagusia bilineata* | Doublelined tonguesole | 1 |
| F-OWF | 88 | *Pentaceros wheeleri* | Slender armorhead | 1 |
| F-OWF | 89 | *Polymixia berndti* | Pacific beardfish | 1 |
| F-OWF | 90 | *Promethichthys prometheus* | Roudi escolar | 1 |
| F-OWF | 91 | *Pseudolabrus eoethinus* | Red naped wrasse | 1 |
| F-OWF | 92 | *Pseudorhombus pentophthalmus* | Fivespot flounder | 1 |
| F-OWF | 93 | *Scopeloberyx robustus* | Paddlenose chimaera | 1 |
| F-OWF | 94 | *Takifugu oblongus* | Lattice blaasop | 1 |
| F-OWF | 95 | *Zanclus cornutus* | Moorish idol | 1 |
| Control | 1 | *Chirocentrus dorab* | Dorab wolf-herring | 728 |
| Control | 2 | *Scatophagus argus* | Spotted scat | 652 |
| Control | 3 | *Plesiops nakaharae* | Nakahara's longfin | 359 |
| Control | 4 | *Heptranchias perlo* | Sharpnose sevengill shark | 309 |
| Control | 5 | *Selenanthias analis* | Pearl-spotted fairy basslet | 301 |
| Control | 6 | *Chimaera phantasma* | Silver chimaera | 285 |
| Control | 7 | *Ventrifossa garmani* | Sagami grenadier | 274 |
| Control | 8 | *Synodus macrops* | Triplecross lizardfish | 259 |
| Control | 9 | *Girella mezina* | Yellowstriped blackfish | 250 |

| | | | | |
|---|---|---|---|---|
| Control | 10 | *Synodus variegatus* | Variegated lizardfish | 242 |
| Control | 11 | *Syngnathus schlegeli* | Seaweed pipefish | 240 |
| Control | 12 | *Epinephelus akaara* | Hong Kong grouper | 210 |
| Control | 13 | *Cephalopholis sonnerati* | Tomato hind | 203 |
| Control | 14 | *Notorynchus cepedianus* | Broadnose sevengill shark | 190 |
| Control | 15 | *Epinephelus merra* | Honeycomb grouper | 186 |
| Control | 16 | *Squalogadus modificatus* | Tadpole whiptail | 177 |
| Control | 17 | *Synaphobranchus kaupii* | Kaup's arrowtooth eel | 169 |
| Control | 18 | *Ablennes hians* | Flat needlefish | 168 |
| Control | 19 | *Polymixia japonica* | Silver eye | 168 |
| Control | 20 | *Muraenichthys gymnopterus* | Anguilliformes Ophichthidae | 161 |
| Control | 21 | *Ophidion muraenolepis* | Blackedge cusk | 159 |
| Control | 22 | *Parapercis ommatura* | Perciformes: Pinguipedidae | 150 |
| Control | 23 | *Hippocampus kuda* | Spotted seahorse | 126 |
| Control | 24 | *Lobotes surinamensis* | Tripletail | 118 |
| Control | 25 | *Synodus oculeus* | Large-eye lizardfish | 115 |
| Control | 26 | *Antennarius striatus* | Striated frogfish | 114 |
| Control | 27 | *Scomber australasicus* | Blue mackerel | 113 |
| Control | 28 | *Amphiprion clarkii* | Yellowtail clownfish | 111 |
| Control | 29 | *Abudefduf sexfasciatus* | Scissortail sergeant | 100 |
| Control | 30 | *Amblyglyphidodon curacao* | Staghorn damselfish | 94 |
| Control | 31 | *Stemonidium hypomelas* | Black serrivomerid eel | 92 |
| Control | 32 | *Oxyconger leptognathus* | Shorttail pike conger | 84 |
| Control | 33 | *Trachurus japonicus* | Japanese jack mackerel | 84 |

| | | | | |
|---|---|---|---|---|
| Control | 34 | *Eurypharynx pelecanoides* | Pelican eel | 79 |
| Control | 35 | *Gymnothorax kidako* | Kidako moray | 77 |
| Control | 36 | *Corythoichthys flavofasciatus* | Network pipefish | 69 |
| Control | 37 | *Sphyraena pinguis* | Red barracuda | 64 |
| Control | 38 | *Euthynnus affinis* | Kawakawa | 63 |
| Control | 39 | *Cynoglossus bilineatus* | Fourlined tonguesole | 60 |
| Control | 40 | *Harriotta raleighana* | Pacific longnose chimaera | 60 |
| Control | 41 | *Valenciennellus tripunctulatus* | Constellationfish | 57 |
| Control | 42 | *Herklotsichthys quadrimaculatus* | Bluestripe herring | 56 |
| Control | 43 | *Albula glossodonta* | Roundjaw bonefish | 53 |
| Control | 44 | *Sigmops gracilis* | Slender fangjaw | 50 |
| Control | 45 | *Fowlerichthys scriptissimus* | Fishing frog | 47 |
| Control | 46 | *Plectroglyphidodon leucozonus* | Singlebar devil | 41 |
| Control | 47 | *Gadus chalcogrammus* | Alaska pollock | 39 |
| Control | 48 | *Cynoglossus joyneri* | Red tonguesole | 33 |
| Control | 49 | *Pempheris schwenkii* | Silver sweeper | 24 |
| Control | 50 | *Pomacentrus moluccensis* | Lemon damsel | 20 |
| Control | 51 | *Bolbometopon muricatum* | Green humphead parrotfish | 17 |
| Control | 52 | *Pomacentrus brachialis* | Charcoal damsel | 16 |
| Control | 53 | *Paraplagusia japonica* | Black cow-tongue | 9 |
| Control | 54 | *Anampses geographicus* | Geographic wrasse | 8 |
| Control | 55 | *Pristipomoides filamentosus* | Crimson jobfish | 8 |
| Control | 56 | *Macropharyngodon meleagris* | Blackspotted wrasse | 7 |
| Control | 57 | *Vinciguerria nimbaria* | Oceanic lightfish | 7 |

| | | | | |
|---|---|---|---|---|
| Control | 58 | *Anguilla marmorata* | Giant mottled eel | 6 |
| Control | 59 | *Anyperodon leucogrammicus* | Slender grouper | 6 |
| Control | 60 | *Girella punctata* | Largescale blackfish | 6 |
| Control | 61 | *Kyphosus bigibbus* | Brown chub | 6 |
| Control | 62 | *Takifugu oblongus* | Lattice blaasop | 6 |
| Control | 63 | *Lophotus capellei* | Unicornfish | 5 |
| Control | 64 | *Thunnus obesus* | Bigeye tuna | 5 |
| Control | 65 | *Chlamydoselachus anguineus* | Frilled shark | 4 |
| Control | 66 | *Paraplagusia bilineata* | Doublelined tonguesole | 4 |
| Control | 67 | *Callogobius okinawae* | Okinawa flap-headed goby | 3 |
| Control | 68 | *Ptereleotris heteroptera* | Blacktail goby | 3 |
| Control | 69 | *Zeus faber* | John dory | 3 |
| Control | 70 | *Alepisaurus ferox* | Long snouted lancetfish | 2 |
| Control | 71 | *Chlorophthalmus albatrossis* | Bigeyed greeneye | 2 |
| Control | 72 | *Coelorinchus longissimus* | - | 2 |
| Control | 73 | *Gerres oyena* | Common silver-biddy | 2 |
| Control | 74 | *Lobianchia gemellarii* | Cocco's lantern fish | 2 |
| Control | 75 | *Lutjanus argentimaculatus* | Mangrove red snapper | 2 |
| Control | 76 | *Nemipterus bathybius* | Yellowbelly threadfin bream | 2 |
| Control | 77 | *Pleurogrammus azonus* | Okhotsk atka mackerel | 2 |
| Control | 78 | *Pterygotrigla ryukyuensis* | Ryukyu gurnard | 2 |
| Control | 79 | *Spratelloides delicatulus* | Delicate round herring | 2 |
| Control | 80 | *Zanclus cornutus* | Moorish idol | 2 |
| Control | 81 | *Acentrogobius suluensis* | Sulu goby | 1 |

| | | | | |
|---|---|---|---|---|
| Control | 82 | *Amblychaeturichthys sciistius* | - | 1 |
| Control | 83 | *Argyropelecus aculeatus* | Lovely hatchetfish | 1 |
| Control | 84 | *Arnoglossus polyspilus* | Many-spotted lefteye flounder | 1 |
| Control | 85 | *Bathygadus antrodes* | - | 1 |
| Control | 86 | *Cheilinus fasciatus* | Redbreasted wrasse | 1 |
| Control | 87 | *Coelorinchus gilberti* | - | 1 |
| Control | 88 | *Coelorinchus hubbsi* | - | 1 |
| Control | 89 | *Dactyloptena peterseni* | Starry flying gurnard | 1 |
| Control | 90 | *Discordipinna griessingeri* | Spikefin goby | 1 |
| Control | 91 | *Dolichopteryx minuscula* | - | 1 |
| Control | 92 | *Engraulis japonicus* | Japanese anchovy | 1 |
| Control | 93 | *Gazza minuta* | Toothpony | 1 |
| Control | 94 | *Gobiodon prolixus* | Elongate coralgoby | 1 |
| Control | 95 | *Hygophum reinhardtii* | Reinhardt's lantern fish | 1 |
| Control | 96 | *Hypoptychus dybowskii* | Korean sandlance | 1 |
| Control | 97 | *Lethrinus atkinsoni* | Pacific yellowtail emperor | 1 |
| Control | 98 | *Lutjanus lutjanus* | Bigeye snapper | 1 |
| Control | 99 | *Metavelifer multiradiatus* | Spinyfin velifer | 1 |
| Control | 100 | *Moringua microchir* | Lesser thrush eel | 1 |
| Control | 101 | *Nemichthys scolopaceus* | Slender snipe eel | 1 |
| Control | 102 | *Neoclinus okazakii* | - | 1 |
| Control | 103 | *Neoditrema ransonnetii* | - | 1 |
| Control | 104 | *Pardachirus pavoninus* | Peacock sole | 1 |
| Control | 105 | *Parupeneus cyclostomus* | Gold-saddle goatfish | 1 |

| | | | | |
|---|---|---|---|---|
| Control | 106 | *Polymixia berndti* | Pacific beardfish | 1 |
| Control | 107 | *Pomacentrus coelestis* | Neon damselfish | 1 |
| Control | 108 | *Poromitra oscitans* | Yawning | 1 |
| Control | 109 | *Pristiophorus japonicus* | Japanese sawshark | 1 |
| Control | 110 | *Promethichthys prometheus* | Roudi escolar | 1 |
| Control | 111 | *Pungitius pungitius* | Ninespine stickleback | 1 |
| Control | 112 | *Rhinochimaera africana* | Paddlenose chimaera | 1 |

Table S4 NASC per mile by transect and depth. NASC per mile ($m^2\ nmi^{-2}$) is presented for each transect and depth layer. Transects FH–F8, F8–FH, S1–S6, and S6–S1 are designated as Navigation A, B, C, and D, respectively. The table also includes the summed NASC across all depth layers for each transect (Sum NASC) and the corresponding transect distance (nmi). Cells with zero values are indicated as NA.

| Sampling Time (Start) | Sampling Time (Finish) | Latitude (Start) | Longitude (Start) | Latitude (Finish) | Longitude (Finish) | Area | Transect | depth (m) | Distance (nmi) | Sum NASC | NASC per mile （m² / nmi²） |
|---|---|---|---|---|---|---|---|---|---|---|---|
| 2025/5/1 14:30 | 2025/5/1 15:21 | 32°40.5678' N | 129°00.3019 ' E | 32°40.4135' N | 128°56.0584 ' E | FOWF | Navigation B | 12.5 | 6 | 0 | NA |
| | | | | | | FOWF | Navigation B | 17.5 | 6 | 0 | NA |
| | | | | | | FOWF | Navigation B | 22.5 | 6 | 46.323754 | 7.72 |
| | | | | | | FOWF | Navigation B | 27.5 | 6 | 76.127366 | 12.69 |
| | | | | | | FOWF | Navigation B | 32.5 | 6 | 139.287323 | 23.21 |
| | | | | | | FOWF | Navigation B | 37.5 | 6 | 220.469025 | 36.74 |
| | | | | | | FOWF | Navigation B | 42.5 | 6 | 224.718302 | 37.45 |
| | | | | | | FOWF | Navigation B | 47.5 | 6 | 188.130359 | 31.36 |
| | | | | | | FOWF | Navigation B | 52.5 | 6 | 249.619862 | 41.60 |
| | | | | | | FOWF | Navigation B | 57.5 | 6 | 268.160122 | 44.69 |
| | | | | | | FOWF | Navigation B | 62.5 | 6 | 272.018283 | 45.34 |
| | | | | | | FOWF | Navigation B | 67.5 | 6 | 166.243481 | 27.71 |
| | | | | | | FOWF | Navigation B | 72.5 | 6 | 121.437576 | 20.24 |
| | | | | | | FOWF | Navigation B | 77.5 | 6 | 78.120335 | 13.02 |
| | | | | | | FOWF | Navigation B | 82.5 | 6 | 0 | NA |
| | | | | | | FOWF | Navigation B | 87.5 | 6 | 0 | NA |
| | | | | | | FOWF | Navigation B | 92.5 | 6 | 0 | NA |

| | | | | | | | | | | | |
|---|---|---|---|---|---|---|---|---|---|---|---|
| | | | | | | FOWF | Navigation B | 97.5 | 6 | 0 | NA |
| | | | | | | FOWF | Navigation B | 102.5 | 6 | 22.502108 | 3.75 |
| | | | | | | FOWF | Navigation B | 107.5 | 6 | 114.873324 | 19.15 |
| | | | | | | FOWF | Navigation B | 112.5 | 6 | 338.144289 | 56.36 |
| | | | | | | FOWF | Navigation B | 117.5 | 6 | 662.346013 | 110.39 |
| | | | | | | FOWF | Navigation B | 122.5 | 6 | 941.766459 | 156.96 |
| | | | | | | FOWF | Navigation B | 127.5 | 6 | 346.308227 | 57.72 |
| | | | | | | FOWF | Navigation B | 132.5 | 6 | 1.808458 | 0.30 |
| 2025/5/1 11:21 | 2025/5/1 12:06 | 32°36.1452' N | 128°56.7681' E | 32°35.7898' N | 129°01.2536' E | Control | Navigation C | 12.5 | 6 | 29.56286 | 4.93 |
| | | | | | | Control | Navigation C | 17.5 | 6 | 76.966789 | 12.83 |
| | | | | | | Control | Navigation C | 22.5 | 6 | 158.822589 | 26.47 |
| | | | | | | Control | Navigation C | 27.5 | 6 | 135.729676 | 22.62 |
| | | | | | | Control | Navigation C | 32.5 | 6 | 160.612805 | 26.77 |
| | | | | | | Control | Navigation C | 37.5 | 6 | 57.896481 | 9.65 |
| | | | | | | Control | Navigation C | 42.5 | 6 | 59.656279 | 9.94 |
| | | | | | | Control | Navigation C | 47.5 | 6 | 52.065631 | 8.68 |
| | | | | | | Control | Navigation C | 52.5 | 6 | 52.363626 | 8.73 |
| | | | | | | Control | Navigation C | 57.5 | 6 | 25.384028 | 4.23 |
| | | | | | | Control | Navigation C | 62.5 | 6 | 27.028913 | 4.50 |

| | | | | | | | | | | | |
|---|---|---|---|---|---|---|---|---|---|---|---|
| | | | | | | Control | Navigation C | 67.5 | 6 | 0 | NA |
| | | | | | | Control | Navigation C | 72.5 | 6 | 0 | NA |
| | | | | | | Control | Navigation C | 77.5 | 6 | 28.713224 | 4.79 |
| | | | | | | Control | Navigation C | 82.5 | 6 | 44.503543 | 7.42 |
| | | | | | | Control | Navigation C | 87.5 | 6 | 0 | NA |
| | | | | | | Control | Navigation C | 92.5 | 6 | 6.756692 | 1.13 |
| | | | | | | Control | Navigation C | 97.5 | 6 | 30.436891 | 5.07 |
| | | | | | | Control | Navigation C | 102.5 | 6 | 78.103472 | 13.02 |
| | | | | | | Control | Navigation C | 107.5 | 6 | 0 | NA |
| | | | | | | Control | Navigation C | 112.5 | 6 | 47.969814 | 7.99 |
| | | | | | | Control | Navigation C | 117.5 | 6 | 25.650752 | 4.28 |
| | | | | | | Control | Navigation C | 122.5 | 6 | 0 | NA |
| | | | | | | Control | Navigation C | 127.5 | 6 | 23.088356 | 3.85 |
| | | | | | | Control | Navigation C | 132.5 | 6 | 0 | NA |
| | | | | | | Control | Navigation C | 137.5 | 6 | 0 | NA |
| | | | | | | Control | Navigation C | 142.5 | 6 | 0 | NA |
| | | | | | | Control | Navigation C | 147.5 | 6 | 78.516337 | 13.09 |
| | | | | | | Control | Navigation C | 152.5 | 6 | 238.065836 | 39.68 |
| | | | | | | Control | Navigation C | 157.5 | 6 | 451.548211 | 75.26 |
| | | | | | | Control | Navigation C | 162.5 | 6 | 607.567703 | 101.26 |

| | | | | | | | | | | | |
|---|---|---|---|---|---|---|---|---|---|---|---|
| | | | | | | Control | Navigation C | 167.5 | 6 | 458.807543 | 76.47 |
| | | | | | | Control | Navigation C | 172.5 | 6 | 569.65905 | 94.94 |
| | | | | | | Control | Navigation C | 177.5 | 6 | 500.705611 | 83.45 |
| | | | | | | Control | Navigation C | 182.5 | 6 | 0.47581 | 0.08 |
| 2025/5/1 12:10 | 2025/5/1 12:44 | 32°35.7830' N | 129°00.5497' E | 32°36.2346' N | 128°36.2364' E | Control | Navigation D | 12.5 | 5.5 | 52.441217 | 8.74 |
| | | | | | | Control | Navigation D | 17.5 | 5.5 | 97.303954 | 16.22 |
| | | | | | | Control | Navigation D | 22.5 | 5.5 | 180.805959 | 30.13 |
| | | | | | | Control | Navigation D | 27.5 | 5.5 | 170.920798 | 28.49 |
| | | | | | | Control | Navigation D | 32.5 | 5.5 | 134.1466 | 22.36 |
| | | | | | | Control | Navigation D | 37.5 | 5.5 | 99.710082 | 16.62 |
| | | | | | | Control | Navigation D | 42.5 | 5.5 | 82.318738 | 13.72 |
| | | | | | | Control | Navigation D | 47.5 | 5.5 | 69.212632 | 11.54 |
| | | | | | | Control | Navigation D | 52.5 | 5.5 | 104.583359 | 17.43 |
| | | | | | | Control | Navigation D | 57.5 | 5.5 | 115.509946 | 19.25 |
| | | | | | | Control | Navigation D | 62.5 | 5.5 | 0 | NA |
| | | | | | | Control | Navigation D | 67.5 | 5.5 | 0 | NA |
| | | | | | | Control | Navigation D | 72.5 | 5.5 | 0 | NA |
| | | | | | | Control | Navigation D | 77.5 | 5.5 | 0 | NA |
| | | | | | | Control | Navigation D | 82.5 | 5.5 | 33.471284 | 5.58 |

| | | | | | | | | | | | |
|---|---|---|---|---|---|---|---|---|---|---|---|
| | | | | | | Control | Navigation D | 87.5 | 5.5 | 0 | NA |
| | | | | | | Control | Navigation D | 92.5 | 5.5 | 66.403566 | 11.07 |
| | | | | | | Control | Navigation D | 97.5 | 5.5 | 0 | NA |
| | | | | | | Control | Navigation D | 102.5 | 5.5 | 35.633576 | 5.94 |
| | | | | | | Control | Navigation D | 107.5 | 5.5 | 43.112617 | 7.19 |
| | | | | | | Control | Navigation D | 112.5 | 5.5 | 25.216362 | 4.20 |
| | | | | | | Control | Navigation D | 117.5 | 5.5 | 48.263865 | 8.04 |
| | | | | | | Control | Navigation D | 122.5 | 5.5 | 0 | NA |
| | | | | | | Control | Navigation D | 127.5 | 5.5 | 23.485864 | 3.91 |
| | | | | | | Control | Navigation D | 132.5 | 5.5 | 0 | NA |
| | | | | | | Control | Navigation D | 137.5 | 5.5 | 22.430011 | 3.74 |
| | | | | | | Control | Navigation D | 142.5 | 5.5 | 55.144153 | 9.19 |
| | | | | | | Control | Navigation D | 147.5 | 5.5 | 25.402624 | 4.23 |
| | | | | | | Control | Navigation D | 152.5 | 5.5 | 119.058564 | 19.84 |
| | | | | | | Control | Navigation D | 157.5 | 5.5 | 404.361083 | 67.39 |
| | | | | | | Control | Navigation D | 162.5 | 5.5 | 776.166838 | 129.36 |
| | | | | | | Control | Navigation D | 167.5 | 5.5 | 408.779564 | 68.13 |
| | | | | | | Control | Navigation D | 172.5 | 5.5 | 417.197677 | 69.53 |
| | | | | | | Control | Navigation D | 177.5 | 5.5 | 626.69654 | 104.45 |
| | | | | | | Control | Navigation D | 182.5 | 5.5 | 56.464278 | 9.41 |

| | | | | | | | | | | | |
|---|---|---|---|---|---|---|---|---|---|---|---|
| 2025/6/5 7:34 | 2025/6/5 8:16 | 32°40.3050' N | 128°56.0480 ' E | 32°40.1081' N | 128°59.9932 ' E | FOWF | Navigation A | 12.5 | 5.5 | 0 | NA |
| | | | | | | FOWF | Navigation A | 17.5 | 5.5 | 26.911179 | 4.89 |
| | | | | | | FOWF | Navigation A | 22.5 | 5.5 | 25.418452 | 4.62 |
| | | | | | | FOWF | Navigation A | 27.5 | 5.5 | 0 | NA |
| | | | | | | FOWF | Navigation A | 32.5 | 5.5 | 165.394784 | 30.07 |
| | | | | | | FOWF | Navigation A | 37.5 | 5.5 | 70.096932 | 12.74 |
| | | | | | | FOWF | Navigation A | 42.5 | 5.5 | 157.527355 | 28.64 |
| | | | | | | FOWF | Navigation A | 47.5 | 5.5 | 55.503161 | 10.09 |
| | | | | | | FOWF | Navigation A | 52.5 | 5.5 | 0 | NA |
| | | | | | | FOWF | Navigation A | 57.5 | 5.5 | 0 | NA |
| | | | | | | FOWF | Navigation A | 62.5 | 5.5 | 0 | NA |
| | | | | | | FOWF | Navigation A | 67.5 | 5.5 | 33.48588 | 6.09 |
| | | | | | | FOWF | Navigation A | 72.5 | 5.5 | 32.550744 | 5.92 |
| | | | | | | FOWF | Navigation A | 77.5 | 5.5 | 28.145353 | 5.12 |
| | | | | | | FOWF | Navigation A | 82.5 | 5.5 | 0 | NA |
| | | | | | | FOWF | Navigation A | 87.5 | 5.5 | 0 | NA |
| | | | | | | FOWF | Navigation A | 92.5 | 5.5 | 2538.403257 | 461.53 |
| | | | | | | FOWF | Navigation A | 97.5 | 5.5 | 326.628994 | 59.39 |
| | | | | | | FOWF | Navigation A | 102.5 | 5.5 | 111.77181 | 20.32 |

| | | | | | | | | | | | |
|---|---|---|---|---|---|---|---|---|---|---|---|
| | | | | | | FOWF | Navigation A | 107.5 | 5.5 | 49.324023 | 8.97 |
| | | | | | | FOWF | Navigation A | 112.5 | 5.5 | 175.066564 | 31.83 |
| | | | | | | FOWF | Navigation A | 117.5 | 5.5 | 0 | NA |
| | | | | | | FOWF | Navigation A | 122.5 | 5.5 | 0 | NA |
| | | | | | | FOWF | Navigation A | 127.5 | 5.5 | 0 | NA |
| | | | | | | FOWF | Navigation A | 132.5 | 5.5 | 0 | NA |
| 2025/6/5 8:16 | 2025/6/5 8:52 | 32°40.1081' N | 128°59.9932' E | 32°40.1906' N | 128°56.0426' E | FOWF | Navigation B | 12.5 | 5 | 0 | NA |
| | | | | | | FOWF | Navigation B | 17.5 | 5 | 0 | NA |
| | | | | | | FOWF | Navigation B | 22.5 | 5 | 96.422177 | 17.53 |
| | | | | | | FOWF | Navigation B | 27.5 | 5 | 0 | NA |
| | | | | | | FOWF | Navigation B | 32.5 | 5 | 66.003062 | 12.00 |
| | | | | | | FOWF | Navigation B | 37.5 | 5 | 121.614357 | 22.11 |
| | | | | | | FOWF | Navigation B | 42.5 | 5 | 36.469034 | 6.63 |
| | | | | | | FOWF | Navigation B | 47.5 | 5 | 29.798531 | 5.42 |
| | | | | | | FOWF | Navigation B | 52.5 | 5 | 0 | NA |
| | | | | | | FOWF | Navigation B | 57.5 | 5 | 21.72817 | 3.95 |
| | | | | | | FOWF | Navigation B | 62.5 | 5 | 70.481978 | 12.81 |
| | | | | | | FOWF | Navigation B | 67.5 | 5 | 40.877581 | 7.43 |
| | | | | | | FOWF | Navigation B | 72.5 | 5 | 0 | NA |

| | | | | | | | | | | | |
|---|---|---|---|---|---|---|---|---|---|---|---|
| | | | | | | FOWF | Navigation B | 77.5 | 5 | 0 | NA |
| | | | | | | FOWF | Navigation B | 82.5 | 5 | 0 | NA |
| | | | | | | FOWF | Navigation B | 87.5 | 5 | 81.184056 | 14.76 |
| | | | | | | FOWF | Navigation B | 92.5 | 5 | 25.069267 | 4.56 |
| | | | | | | FOWF | Navigation B | 97.5 | 5 | 0 | NA |
| | | | | | | FOWF | Navigation B | 102.5 | 5 | 0 | NA |
| | | | | | | FOWF | Navigation B | 107.5 | 5 | 0 | NA |
| | | | | | | FOWF | Navigation B | 112.5 | 5 | 238.732212 | 43.41 |
| | | | | | | FOWF | Navigation B | 117.5 | 5 | 73.866436 | 13.43 |
| | | | | | | FOWF | Navigation B | 122.5 | 5 | 24.017098 | 4.37 |
| | | | | | | FOWF | Navigation B | 127.5 | 5 | 0.469636 | 0.09 |
| 2025/6/5 10:17 | 2025/6/5 10:49 | 32°35.9753' N | 129°00.4695 ' E | 32°35.9432' N | 128°57.0869 ' E | Control | Navigation C | 12.5 | 4.5 | 0 | NA |
| | | | | | | Control | Navigation C | 17.5 | 4.5 | 0 | NA |
| | | | | | | Control | Navigation C | 22.5 | 4.5 | 0 | NA |
| | | | | | | Control | Navigation C | 27.5 | 4.5 | 0 | NA |
| | | | | | | Control | Navigation C | 32.5 | 4.5 | 0 | NA |
| | | | | | | Control | Navigation C | 37.5 | 4.5 | 0 | NA |
| | | | | | | Control | Navigation C | 42.5 | 4.5 | 0 | NA |
| | | | | | | Control | Navigation C | 47.5 | 4.5 | 0 | NA |

| | | | | | | | | | | | |
|---|---|---|---|---|---|---|---|---|---|---|---|
| | | | | | | Control | Navigation C | 52.5 | 4.5 | 0 | NA |
| | | | | | | Control | Navigation C | 57.5 | 4.5 | 0 | NA |
| | | | | | | Control | Navigation C | 62.5 | 4.5 | 0 | NA |
| | | | | | | Control | Navigation C | 67.5 | 4.5 | 0 | NA |
| | | | | | | Control | Navigation C | 72.5 | 4.5 | 0 | NA |
| | | | | | | Control | Navigation C | 77.5 | 4.5 | 0 | NA |
| | | | | | | Control | Navigation C | 82.5 | 4.5 | 0 | NA |
| | | | | | | Control | Navigation C | 87.5 | 4.5 | 0 | NA |
| | | | | | | Control | Navigation C | 92.5 | 4.5 | 0 | NA |
| | | | | | | Control | Navigation C | 97.5 | 4.5 | 0 | NA |
| | | | | | | Control | Navigation C | 102.5 | 4.5 | 12.430138 | 2.26 |
| | | | | | | Control | Navigation C | 107.5 | 4.5 | 51.711921 | 9.40 |
| | | | | | | Control | Navigation C | 112.5 | 4.5 | 12.401849 | 2.25 |
| | | | | | | Control | Navigation C | 117.5 | 4.5 | 0 | NA |
| | | | | | | Control | Navigation C | 122.5 | 4.5 | 0 | NA |
| | | | | | | Control | Navigation C | 127.5 | 4.5 | 0 | NA |
| | | | | | | Control | Navigation C | 132.5 | 4.5 | 2.379926 | 0.43 |
| | | | | | | Control | Navigation C | 137.5 | 4.5 | 9.207311 | 1.67 |
| | | | | | | Control | Navigation C | 142.5 | 4.5 | 33.547054 | 6.10 |
| | | | | | | Control | Navigation C | 147.5 | 4.5 | 59.019608 | 10.73 |

| | | | | | | | | | | | |
|---|---|---|---|---|---|---|---|---|---|---|---|
| | | | | | | Control | Navigation C | 152.5 | 4.5 | 233.964348 | 42.54 |
| | | | | | | Control | Navigation C | 157.5 | 4.5 | 367.724531 | 66.86 |
| | | | | | | Control | Navigation C | 162.5 | 4.5 | 179.342948 | 32.61 |
| | | | | | | Control | Navigation C | 167.5 | 4.5 | 179.770595 | 32.69 |
| | | | | | | Control | Navigation C | 172.5 | 4.5 | 13.64534 | 2.48 |
| 2025/6/5 10:49 | 2025/6/5 11:24 | 32°35.8461' N | 128°57.1030 ' E | 32°35.9549' N | 129°00.4735 ' E | Control | Navigation D | 12.5 | 4.5 | 0 | NA |
| | | | | | | Control | Navigation D | 17.5 | 4.5 | 0 | NA |
| | | | | | | Control | Navigation D | 22.5 | 4.5 | 0 | NA |
| | | | | | | Control | Navigation D | 27.5 | 4.5 | 0 | NA |
| | | | | | | Control | Navigation D | 32.5 | 4.5 | 0 | NA |
| | | | | | | Control | Navigation D | 37.5 | 4.5 | 0 | NA |
| | | | | | | Control | Navigation D | 42.5 | 4.5 | 0 | NA |
| | | | | | | Control | Navigation D | 47.5 | 4.5 | 0 | NA |
| | | | | | | Control | Navigation D | 52.5 | 4.5 | 0 | NA |
| | | | | | | Control | Navigation D | 57.5 | 4.5 | 0 | NA |
| | | | | | | Control | Navigation D | 62.5 | 4.5 | 0 | NA |
| | | | | | | Control | Navigation D | 67.5 | 4.5 | 0 | NA |
| | | | | | | Control | Navigation D | 72.5 | 4.5 | 0 | NA |
| | | | | | | Control | Navigation D | 77.5 | 4.5 | 0 | NA |

| | | | | | | | | | | | |
|---|---|---|---|---|---|---|---|---|---|---|---|
| | | | | | | Control | Navigation D | 82.5 | 4.5 | 0 | NA |
| | | | | | | Control | Navigation D | 87.5 | 4.5 | 0 | NA |
| | | | | | | Control | Navigation D | 92.5 | 4.5 | 0 | NA |
| | | | | | | Control | Navigation D | 97.5 | 4.5 | 0 | NA |
| | | | | | | Control | Navigation D | 102.5 | 4.5 | 0 | NA |
| | | | | | | Control | Navigation D | 107.5 | 4.5 | 0 | NA |
| | | | | | | Control | Navigation D | 112.5 | 4.5 | 27.97275 | 6.22 |
| | | | | | | Control | Navigation D | 117.5 | 4.5 | 73.758803 | 16.39 |
| | | | | | | Control | Navigation D | 122.5 | 4.5 | 11.426537 | 2.54 |
| | | | | | | Control | Navigation D | 127.5 | 4.5 | 17.157067 | 3.81 |
| | | | | | | Control | Navigation D | 132.5 | 4.5 | 8.751739 | 1.94 |
| | | | | | | Control | Navigation D | 137.5 | 4.5 | 0 | NA |
| | | | | | | Control | Navigation D | 142.5 | 4.5 | 116.43004 | 25.87 |
| | | | | | | Control | Navigation D | 147.5 | 4.5 | 88.580052 | 19.68 |
| | | | | | | Control | Navigation D | 152.5 | 4.5 | 382.622538 | 85.03 |
| | | | | | | Control | Navigation D | 157.5 | 4.5 | 499.633846 | 111.03 |
| | | | | | | Control | Navigation D | 162.5 | 4.5 | 337.253466 | 74.95 |
| | | | | | | Control | Navigation D | 167.5 | 4.5 | 105.396516 | 23.42 |
| | | | | | | Control | Navigation D | 172.5 | 4.5 | 71.452333 | 15.88 |
| 2025/8/20 | 2025/8/20 | 32°40.323' | 128°56.076' | 32°40.485' | 129°00.514' | FOWF | Navigation A | 12.5 | 6 | 36.512042 | 5.62 |

| | | | | | | | | | | | |
|---|---|---|---|---|---|---|---|---|---|---|---|
| 17:39 | 18:27 | N | E | N | E | | | | | | |
| | | | | | | FOWF | Navigation A | 17.5 | 6 | 0 | NA |
| | | | | | | FOWF | Navigation A | 22.5 | 6 | 0 | NA |
| | | | | | | FOWF | Navigation A | 27.5 | 6 | 0 | NA |
| | | | | | | FOWF | Navigation A | 32.5 | 6 | 28.99258 | 4.46 |
| | | | | | | FOWF | Navigation A | 37.5 | 6 | 32.716946 | 5.03 |
| | | | | | | FOWF | Navigation A | 42.5 | 6 | 24.70325 | 3.80 |
| | | | | | | FOWF | Navigation A | 47.5 | 6 | 0 | NA |
| | | | | | | FOWF | Navigation A | 52.5 | 6 | 0 | NA |
| | | | | | | FOWF | Navigation A | 57.5 | 6 | 0 | NA |
| | | | | | | FOWF | Navigation A | 62.5 | 6 | 0 | NA |
| | | | | | | FOWF | Navigation A | 67.5 | 6 | 49.177297 | 7.57 |
| | | | | | | FOWF | Navigation A | 72.5 | 6 | 64.633516 | 9.94 |
| | | | | | | FOWF | Navigation A | 77.5 | 6 | 41.517525 | 6.39 |
| | | | | | | FOWF | Navigation A | 82.5 | 6 | 22.193865 | 3.41 |
| | | | | | | FOWF | Navigation A | 87.5 | 6 | 35.583288 | 5.47 |
| | | | | | | FOWF | Navigation A | 92.5 | 6 | 119.286225 | 18.35 |
| | | | | | | FOWF | Navigation A | 97.5 | 6 | 109.010139 | 16.77 |
| | | | | | | FOWF | Navigation A | 102.5 | 6 | 989.377543 | 152.21 |
| | | | | | | FOWF | Navigation A | 107.5 | 6 | 0 | NA |

| | | | | | | | | | | | |
|---|---|---|---|---|---|---|---|---|---|---|---|
| | | | | | | FOWF | Navigation A | 112.5 | 6 | 0 | NA |
| | | | | | | FOWF | Navigation A | 117.5 | 6 | 0 | NA |
| | | | | | | FOWF | Navigation A | 122.5 | 6 | 24.478457 | 3.77 |
| | | | | | | FOWF | Navigation A | 127.5 | 6 | 9.892733 | 1.52 |
| | | | | | | FOWF | Navigation A | 132.5 | 6 | 0.748997 | 0.12 |
| 2025/8/20 16:51 | 2025/8/20 17:39 | 32°40.279' N | 129°00.388' E | 32°40.323' N | 128°56.076' E | FOWF | Navigation B | 12.5 | 6.5 | 22.337432 | 3.44 |
| | | | | | | FOWF | Navigation B | 17.5 | 6.5 | 0 | NA |
| | | | | | | FOWF | Navigation B | 22.5 | 6.5 | 0 | NA |
| | | | | | | FOWF | Navigation B | 27.5 | 6.5 | 0 | NA |
| | | | | | | FOWF | Navigation B | 32.5 | 6.5 | 0 | NA |
| | | | | | | FOWF | Navigation B | 37.5 | 6.5 | 0 | NA |
| | | | | | | FOWF | Navigation B | 42.5 | 6.5 | 0 | NA |
| | | | | | | FOWF | Navigation B | 47.5 | 6.5 | 0 | NA |
| | | | | | | FOWF | Navigation B | 52.5 | 6.5 | 0 | NA |
| | | | | | | FOWF | Navigation B | 57.5 | 6.5 | 0 | NA |
| | | | | | | FOWF | Navigation B | 62.5 | 6.5 | 0 | NA |
| | | | | | | FOWF | Navigation B | 67.5 | 6.5 | 0 | NA |
| | | | | | | FOWF | Navigation B | 72.5 | 6.5 | 0 | NA |
| | | | | | | FOWF | Navigation B | 77.5 | 6.5 | 0 | NA |

| | | | | | | | | | | | |
|---|---|---|---|---|---|---|---|---|---|---|---|
| | | | | | | FOWF | Navigation B | 82.5 | 6.5 | 0 | NA |
| | | | | | | FOWF | Navigation B | 87.5 | 6.5 | 0 | NA |
| | | | | | | FOWF | Navigation B | 92.5 | 6.5 | 0 | NA |
| | | | | | | FOWF | Navigation B | 97.5 | 6.5 | 0 | NA |
| | | | | | | FOWF | Navigation B | 102.5 | 6.5 | 0 | NA |
| | | | | | | FOWF | Navigation B | 107.5 | 6.5 | 0 | NA |
| | | | | | | FOWF | Navigation B | 112.5 | 6.5 | 218.920445 | 33.68 |
| | | | | | | FOWF | Navigation B | 117.5 | 6.5 | 258.359669 | 39.75 |
| | | | | | | FOWF | Navigation B | 122.5 | 6.5 | 0 | NA |
| | | | | | | FOWF | Navigation B | 127.5 | 6.5 | 0 | NA |
| | | | | | | FOWF | Navigation B | 132.5 | 6.5 | 17.651954 | 2.72 |
| 2025/8/20 14:10 | 2025/8/20 14:44 | 32°35.897' N | 128°57.031' E | 32°35.898' N | 129°00.292' E | Control | Navigation C | 12.5 | 4.5 | 32.344605 | 8.09 |
| | | | | | | Control | Navigation C | 17.5 | 4.5 | 21.825241 | 5.46 |
| | | | | | | Control | Navigation C | 22.5 | 4.5 | 0 | NA |
| | | | | | | Control | Navigation C | 27.5 | 4.5 | 0 | NA |
| | | | | | | Control | Navigation C | 32.5 | 4.5 | 0 | NA |
| | | | | | | Control | Navigation C | 37.5 | 4.5 | 0 | NA |
| | | | | | | Control | Navigation C | 42.5 | 4.5 | 0 | NA |
| | | | | | | Control | Navigation C | 47.5 | 4.5 | 0 | NA |

| | | | | | | | | | | | |
|---|---|---|---|---|---|---|---|---|---|---|---|
| | | | | | | Control | Navigation C | 52.5 | 4.5 | 0 | NA |
| | | | | | | Control | Navigation C | 57.5 | 4.5 | 0 | NA |
| | | | | | | Control | Navigation C | 62.5 | 4.5 | 0 | NA |
| | | | | | | Control | Navigation C | 67.5 | 4.5 | 0 | NA |
| | | | | | | Control | Navigation C | 72.5 | 4.5 | 0 | NA |
| | | | | | | Control | Navigation C | 77.5 | 4.5 | 0 | NA |
| | | | | | | Control | Navigation C | 82.5 | 4.5 | 0 | NA |
| | | | | | | Control | Navigation C | 87.5 | 4.5 | 0 | NA |
| | | | | | | Control | Navigation C | 92.5 | 4.5 | 0 | NA |
| | | | | | | Control | Navigation C | 97.5 | 4.5 | 0 | NA |
| | | | | | | Control | Navigation C | 102.5 | 4.5 | 0 | NA |
| | | | | | | Control | Navigation C | 107.5 | 4.5 | 0 | NA |
| | | | | | | Control | Navigation C | 112.5 | 4.5 | 0 | NA |
| | | | | | | Control | Navigation C | 117.5 | 4.5 | 0 | NA |
| | | | | | | Control | Navigation C | 122.5 | 4.5 | 0 | NA |
| | | | | | | Control | Navigation C | 127.5 | 4.5 | 0 | NA |
| | | | | | | Control | Navigation C | 132.5 | 4.5 | 0 | NA |
| | | | | | | Control | Navigation C | 137.5 | 4.5 | 0 | NA |
| | | | | | | Control | Navigation C | 142.5 | 4.5 | 0 | NA |
| | | | | | | Control | Navigation C | 147.5 | 4.5 | 0 | NA |

| | | | | | | | | | | | |
|---|---|---|---|---|---|---|---|---|---|---|---|
| | | | | | | Control | Navigation C | 152.5 | 4.5 | 0 | NA |
| | | | | | | Control | Navigation C | 157.5 | 4.5 | 0 | NA |
| | | | | | | Control | Navigation C | 162.5 | 4.5 | 0 | NA |
| | | | | | | Control | Navigation C | 167.5 | 4.5 | 162.053663 | 40.51 |
| | | | | | | Control | Navigation C | 172.5 | 4.5 | 0 | NA |
| | | | | | | Control | Navigation C | 177.5 | 4.5 | 0 | NA |
| 2025/8/20 14:44 | 2025/8/20 15:14 | 32°35.898' N | 129°00.292' E | 32°35.836' N | 128°57.049' E | Control | Navigation D | 12.5 | 4 | 0 | NA |
| | | | | | | Control | Navigation D | 17.5 | 4 | 22.838522 | 5.08 |
| | | | | | | Control | Navigation D | 22.5 | 4 | 0 | NA |
| | | | | | | Control | Navigation D | 27.5 | 4 | 0 | NA |
| | | | | | | Control | Navigation D | 32.5 | 4 | 0 | NA |
| | | | | | | Control | Navigation D | 37.5 | 4 | 0 | NA |
| | | | | | | Control | Navigation D | 42.5 | 4 | 0 | NA |
| | | | | | | Control | Navigation D | 47.5 | 4 | 0 | NA |
| | | | | | | Control | Navigation D | 52.5 | 4 | 0 | NA |
| | | | | | | Control | Navigation D | 57.5 | 4 | 0 | NA |
| | | | | | | Control | Navigation D | 62.5 | 4 | 0 | NA |
| | | | | | | Control | Navigation D | 67.5 | 4 | 48.551329 | 10.79 |
| | | | | | | Control | Navigation D | 72.5 | 4 | 79.807144 | 17.73 |

| | | | | | | | | | | | |
|---|---|---|---|---|---|---|---|---|---|---|---|
| | | | | | | Control | Navigation D | 77.5 | 4 | 0 | NA |
| | | | | | | Control | Navigation D | 82.5 | 4 | 0 | NA |
| | | | | | | Control | Navigation D | 87.5 | 4 | 0 | NA |
| | | | | | | Control | Navigation D | 92.5 | 4 | 0 | NA |
| | | | | | | Control | Navigation D | 97.5 | 4 | 0 | NA |
| | | | | | | Control | Navigation D | 102.5 | 4 | 0 | NA |
| | | | | | | Control | Navigation D | 107.5 | 4 | 0 | NA |
| | | | | | | Control | Navigation D | 112.5 | 4 | 0 | NA |
| | | | | | | Control | Navigation D | 117.5 | 4 | 0 | NA |
| | | | | | | Control | Navigation D | 122.5 | 4 | 0 | NA |
| | | | | | | Control | Navigation D | 127.5 | 4 | 0 | NA |
| | | | | | | Control | Navigation D | 132.5 | 4 | 0 | NA |
| | | | | | | Control | Navigation D | 137.5 | 4 | 0 | NA |
| | | | | | | Control | Navigation D | 142.5 | 4 | 0 | NA |
| | | | | | | Control | Navigation D | 147.5 | 4 | 0 | NA |
| | | | | | | Control | Navigation D | 152.5 | 4 | 0 | NA |
| | | | | | | Control | Navigation D | 157.5 | 4 | 0 | NA |
| | | | | | | Control | Navigation D | 162.5 | 4 | 0 | NA |
| | | | | | | Control | Navigation D | 167.5 | 4 | 0 | NA |
| | | | | | | Control | Navigation D | 172.5 | 4 | 0 | NA |

| | | | | | | | | | | | |
|---|---|---|---|---|---|---|---|---|---|---|---|
| | | | | | | Control | Navigation D | 177.5 | 4 | 0 | NA |

Fig. S1 Monthly echograms for each slalom transect obtained using a scientific echosounder. Acoustic data were collected using a scientific echosounder (EK80; Simrad Kongsberg Maritime AS, Horten, Norway) operated in continuous-wave (CW) mode with a transmit power of 150 W, frequency of 200 kHz, pulse duration of 1.024 ms, beam width of 7°, and ping interval of 1000 ms. Echograms were visualized using Echoview software (version 9.0; Echoview Software Pty Ltd). Colors represent volume backscattering strength (Sv) displayed using the standard color palette. The horizontal green line in the mid-water layer indicates the draft depth of the wind turbine structure (76 m).

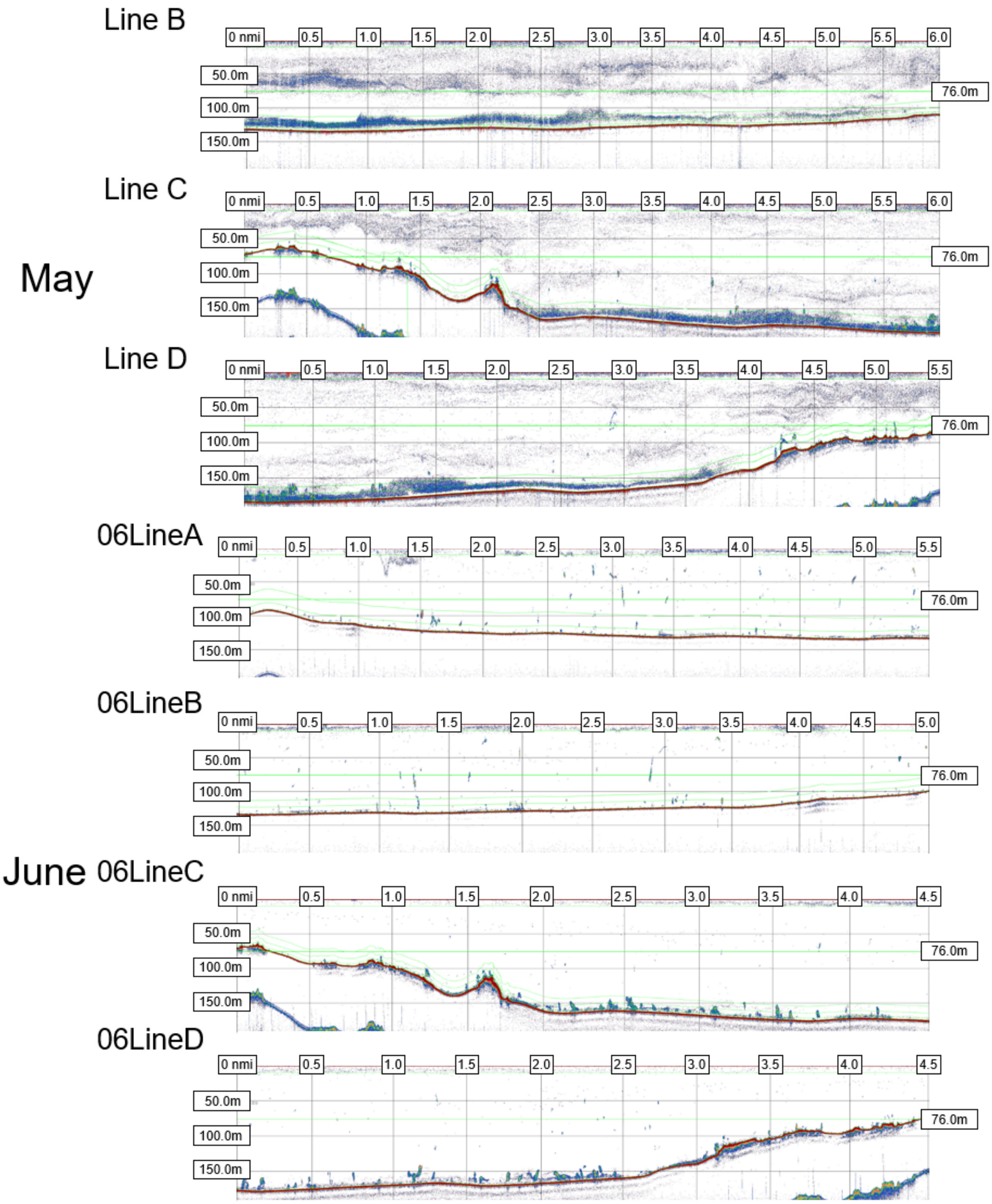

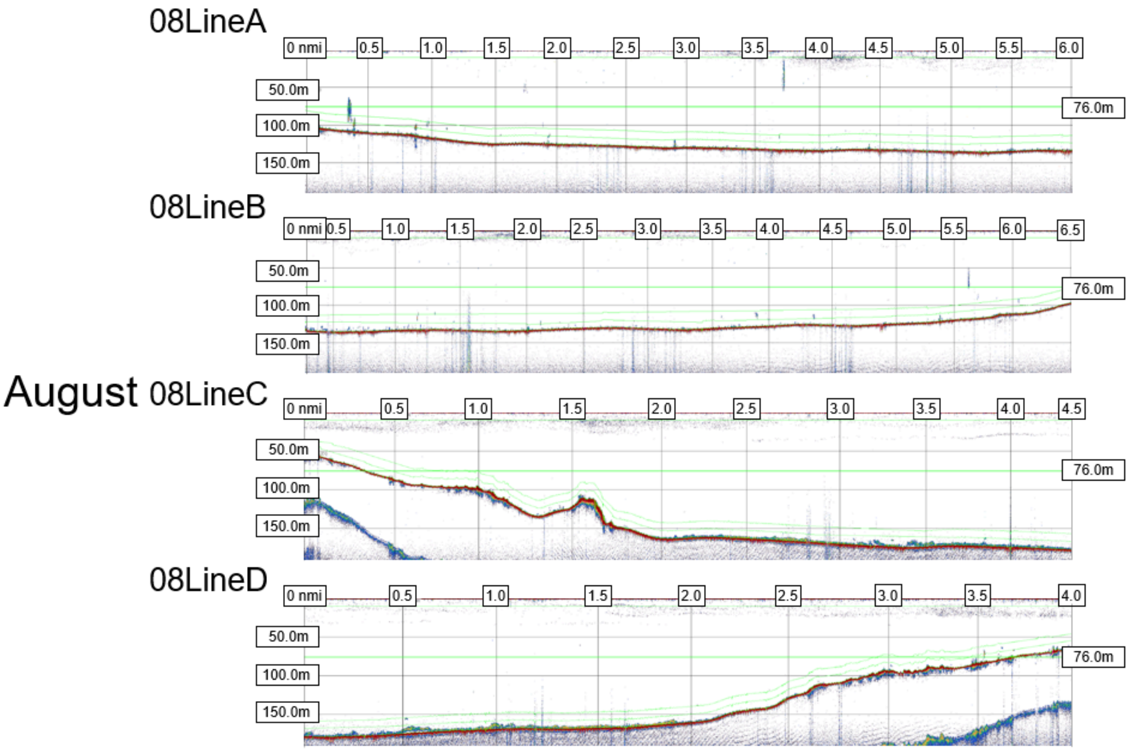

August
08LineA
0 nmi
0.5
1.0
1.5
2.0
2.5
3.0
3.5
4.0
4.5
5.0
5.5
6.0
50.0m
100.0m
150.0m
76.0m
08LineB
0 nmi
0.5
1.0
1.5
2.0
2.5
3.0
3.5
4.0
4.5
5.0
5.5
6.0
6.5
50.0m
100.0m
150.0m
76.0m
08LineC
0 nmi
0.5
1.0
1.5
2.0
2.5
3.0
3.5
4.0
4.5
50.0m
100.0m
150.0m
76.0m
08LineD
0 nmi
0.5
1.0
1.5
2.0
2.5
3.0
3.5
4.0
50.0m
100.0m
150.0m
76.0m
